# Epitranscriptomic control of stress adaptations in *Escherichia coli*

**DOI:** 10.1101/2025.10.14.682255

**Authors:** Sebastián Riquelme-Barrios, Siobhan A. Cusack, Luis Rivera-Montero, Leonardo Vásquez-Camus, Korinna Burdack, Sophie Brameyer, Maximilian Berg, G. Nur Yeşiltaç-Tosun, Stefanie Kaiser, Pascal Giehr, Kirsten Jung

## Abstract

Bacterial stress responses have been studied at the phenotypic, transcriptional, and translational levels, demonstrating the presence of an “alarm” phase immediately after stress exposure. However, the contributions of RNA modifications during stress adaptation remain largely unexplored. Here, we map the epitranscriptomic changes of *Escherichia coli* after exposure to oxidative and acid stress using direct RNA sequencing of mRNA, rRNA, and tRNA, combined with mass spectrometry, deletion mutant phenotyping, and single-nucleotide PCR. We identified widespread, dynamic RNA modifications that include central metabolism transcripts and increased levels of rRNA methylations (m^4^Cm and m^5^C) under both stresses, with potential consequences for translation. In uncharged tRNAs, stress-specific modifications via the Mnm and Q pathways accumulated at the wobble position; these modifications proved crucial for survival. Together, these findings reveal a multifaceted layer of post-transcriptional regulation, establishing the first comprehensive view of the bacterial epitranscriptome during the alarm phase of stress adaptation.

## INTRODUCTION

Bacteria are often subjected to environmental stresses, including sudden changes in pH, temperature, osmolarity, nutrient availability, or reactive oxygen species levels. Bacterial responses are triggered by stress sensing, during which adverse external conditions are identified and transduced via signaling processes. The earliest phase of the *E. coli* stress response after sensing can be thought of as the stress “alarm” phase (according to the biological stress concept^1^), during which signaling cascades are activated to trigger gene expression and growth is halted to redirect resources toward recovery (**Fig. 1a**).

**Figure 1.**
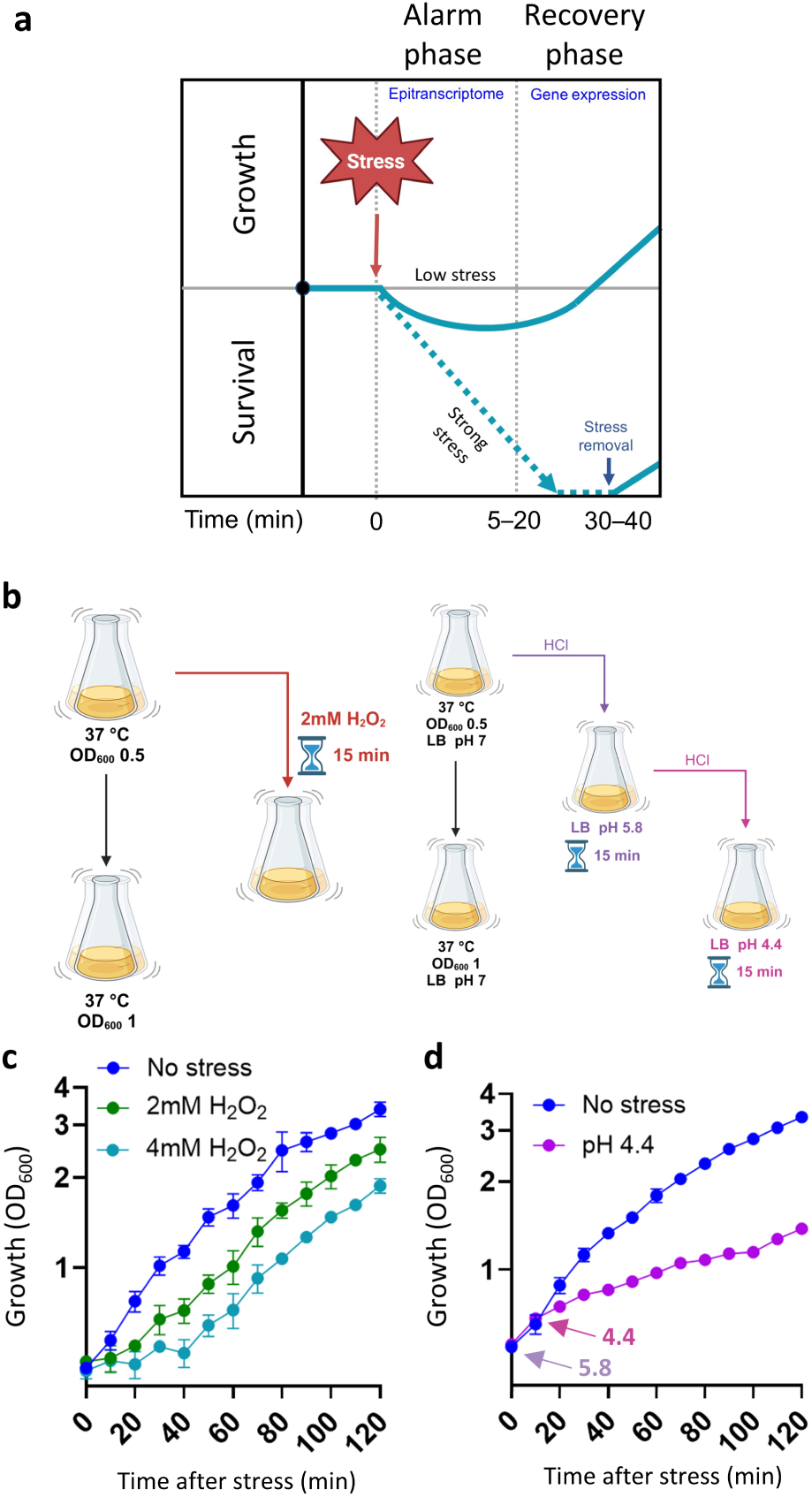
The alarm phase in *E. coli* is characterized by decreased and delayed growth. a, Illustration of the bacterial stress response. The first stage of the stress response is the alarm phase, during which an adverse condition is sensed and growth slows or halts. In the case of a relatively mild stress, bacteria proceed to the recovery phase, during which a new set of genes is expressed and growth resumes. Under strong stress, the bacteria die or survive in a dormant state, from which they can continue to grow once the stimulus is removed. **b,** Overview of the oxidative and acid stress protocols used in the present study. For each stress condition, two flasks of *E. coli* were grown in parallel at 37 °C to an OD_600_ of 0.5. For oxidative stress, H_2_O_2_ was added to one flask to a final concentration of 2 mM and cultivation was continued at 37 °C for 15 min. For acid stress, HCl was added to one flask to decrease the pH to 5.8. Cultivation was then continued at 37 °C for 15 min, after which additional HCl was added to reach pH 4.4. For both stress conditions, the untreated flask remained incubating at 37 °C. Cells were then harvested and RNA was extracted. **c**, Growth of control (no stress) *E. coli* and those exposed to 2 or 4 mM H_2_O_2_. **d**, Growth of control (no stress) *E. coli* and those grown in pH 4.4 media.

Modifications of bacterial tRNAs and rRNAs have been characterized in detail, with roles in structure, catalytic activity, assembly, maturation, translation fidelity, tRNA structure, and codon recognition^2–8^. *In vitro* studies have shown that modifications in mRNAs can impair codon–anticodon interactions, destabilize the ribosome, and increase amino acid substitution errors, impacting reading accuracy at stop codons^9–11^. Previous work from our group demonstrated RNA modifications in *E. coli* that vary between growth stages^12^. More recently, our group identified putative modification sites in *E. coli* mRNAs that exhibit altered abundance in response to heat stress^13^.

The roles of some epitranscriptomic marks during bacterial stress responses have been characterized, particularly in rRNAs. For example, C1402 of the decoding center in the 16S rRNA is modified by both RsmI (which forms Cm) and RsmH^14^ (m^4^C), together forming *N*^4^,2′-*O*-dimethylcytidine (m^4^Cm); these enzymes are implicated in virulence and oxidative stress resistance in *Staphylococcus aureus*^15^. Levels of 5-hydroxycytidine (ho^5^C) at C2501 of the *E. coli* 23S rRNA increase during oxidative stress, impairing protein synthesis and thus conferring a protective effect^16^.

In addition to the rRNA, the roles of some bacterial tRNA modifications throughout the stress response have been defined. In *Pseudomonas aeruginosa*, tRNA methylation (by members of the Trm family) is necessary for resistance to oxidative stress and is involved in survival during infection. Specifically, the absence of *trmJ* increases sensitivity to H_2_O_2_^17^, and Δ*trmB* mutants have impaired expression of mRNAs enriched in Phe and Asp codons, such as *katB* and *katA*^18^. Similarly, mutants for *ttcA*, which catalyzes post-transcriptional thiolation of C32 (s^2^C32), show H_2_O_2_ hypersensitivity and attenuated infection capacity^19^. Overall, H_2_O_2_ exposure produces an increase in 7-methylguanosine (m^7^G) levels; relatedly, a lack of functional *trmB* produces an oxidative stress-sensitive phenotype in *P. aeruginosa*^18^. In response to acid stress, one *E. coli* screen identified the tRNA modification gene *mnmE* as important for growth^20^. A *Streptococcus mutans* mutant lacking *gidA* (a homolog of the *E. coli* modification gene *mnmG*, which together with *mnmE* is responsible for 5-methylaminomethyl-2-thiouridine [mnm^5^s^2^U] modifications) shows increased sensitivity to mild acid stress. Δ*mnmG* and Δ*mnmE* mutants also have difficulties adapting to temperature and osmotic stresses^21^. A similar trend was observed in *Cronobacter sakazakii,* where transposon mutagenesis of *mnmG* causes increased acid sensitivity^22^.

Some stress conditions alter modifications in the wobble position of the tRNA anticodon specifically. These modifications change the binding preference of the anticodon, promoting non-Watson–Crick interactions with the corresponding codon. In some cases, codons that require a modification in the wobble position for translation (i.e., modification-dependent codons) are more abundant among mRNAs that are expressed in response to stress or specific metabolic signals. Conditions that increase wobble-position modification levels can thus initiate translation of transcripts containing a high abundance of modification-dependent codons^23^ in a regulatory mechanism referred to as modification tunable transcripts (MoTTs)^24^. Although some instances of MoTTs regulation have been described in *E. coli*, few stress-responsive MoTTs have been identified in this organism^25^.

Despite the numerous insights yielded by prior studies of tRNA and rRNA modifications under stress conditions, a comprehensive analysis of the *E. coli* epitranscriptomic response to oxidative or acid stress during the stress alarm phase has not yet been conducted. To uncover specific and general epitranscriptomic responses to stress conditions during this phase, we conducted a multifaceted analysis incorporating direct RNA sequencing (DRS), mass spectrometry (MS), single-nucleotide PCR, and mutant phenotype characterization. This allowed us to assess changes in the abundance of mRNA, tRNA, and rRNA modifications in response to stress, revealing both specific and unique stress responses.

## RESULTS

### Nanopore sequencing uncovers stress-responsive epitranscriptomic signatures of mRNAs with central functions

In the present study, we sought to measure the bacterial stress response from an epitranscriptomic perspective, focusing on the alarm phase during oxidative-stress (H_2_O_2_) and acid-stress (HCl) conditions (**Fig. 1a, b**). Oxidative stress was characterized by a dose-dependent lag phase of ∼20 and ∼35 min at 2 and 4 mM H_2_O_2_, respectively, although the growth rate remained unaffected (**Fig. 1c**). Under acid stress, we observed a decrease in the growth rate (**Fig. 1d**). These periods of decreased or halted growth corresponded to the alarm phase of the stress response (**Fig. 1a**).

We next implemented DRS following previously established protocols^13,26,27^ to analyze epitranscriptomic changes during the alarm phase in no-stress controls and samples exposed to oxidative- or acid-stress conditions. A previously developed method^13^ enabled simultaneous capture of rRNAs, mRNAs, ncRNAs, and uncharged tRNAs in a single mRNA-enriched (mRNAe) sample (**Fig. 2a; Table S1)**. Additionally, we sequenced modification-free *in vitro* transcribed (IVT) RNA to control for intrinsic and sequence-specific errors (**Fig. 2a**). Analysis of the sequencing reads demonstrated high depth of coverage **(Extended Data Fig. 1a, b)**, the expected CDS enrichment **(Extended Data Fig. 1c)**, and transcriptome-level stress responses comparable to those described in previous reports^28,29^ **(Extended Data Fig. 1d–g)**.

**Figure 2.**
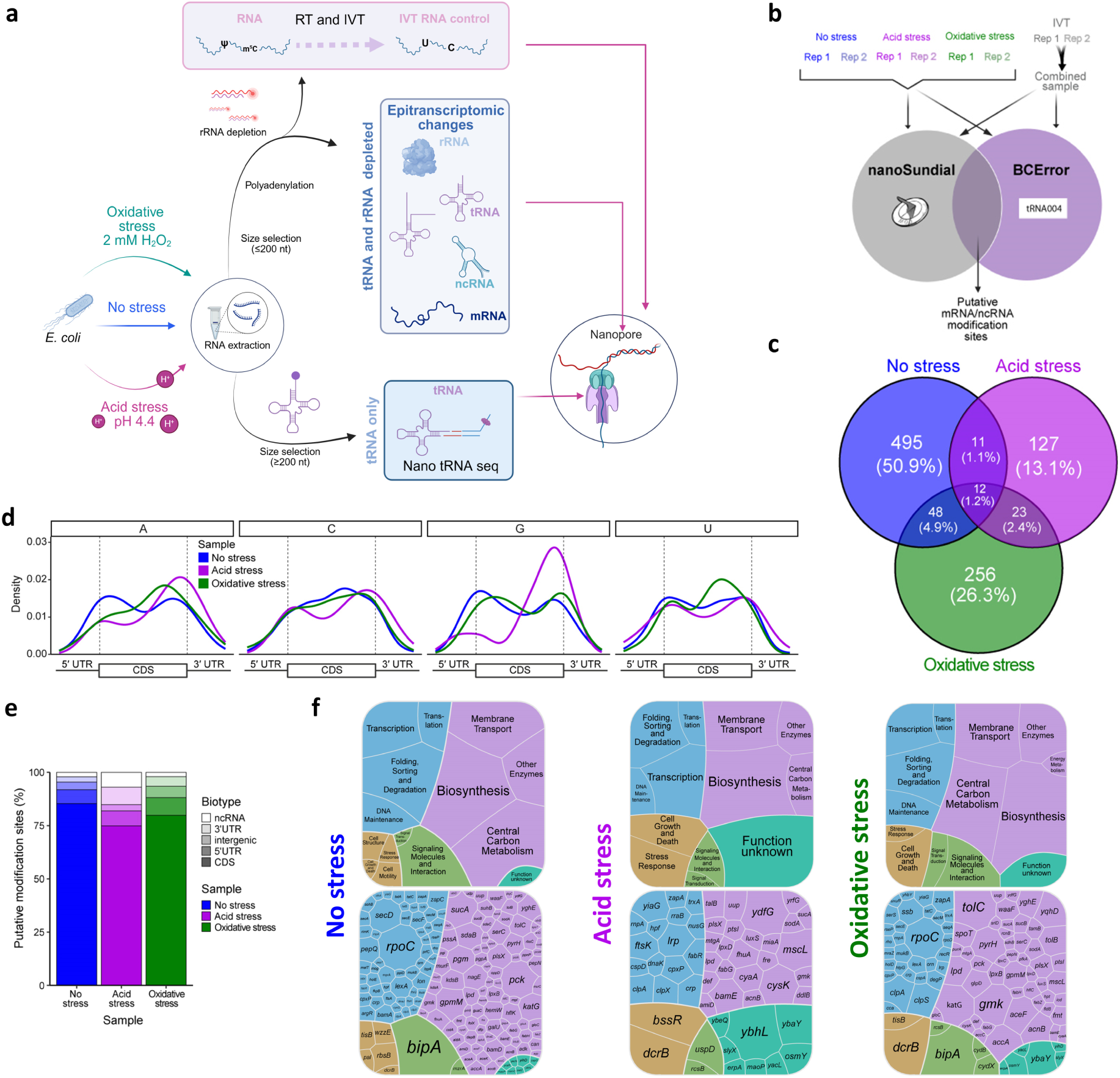
Stress-responsive epitranscriptomic signatures in mRNAs. a, Schematic of the experimental design: sample treatment, library preparation, and Nanopore sequencing of RNA from control and stressed *E. coli* and production of an *in vitro* transcribed (IVT) control. **b,** Graphical illustration of the putative novel modification detection approach. RNA was extracted from two biological replicate samples of *E. coli* grown under acid-stress, oxidative-stress, and no-stress conditions. An unmodified IVT control was generated from the RNA of no-stress samples. After direct RNA sequencing (DRS), putative modification sites outside the tRNA and rRNA were detected using basecalling error (BCError) as calculated with a publicly available script (https://github.com/rnabioco/tRNA004/blob/main/src/modifications/get_bcerror_freqs.py) and a previously published modification-detection program, nanoSundial. Modification sites/regions were considered in further analyses if they were supported by both programs in both biological replicate samples. **c,** Unique and shared putative modification sites/regions between treatment groups. **d,** Distribution of putative mRNA modifications across the 5′ untranslated region (UTR), coding sequence (CDS), and 3′ UTR in each sample type. **e,** Biotype distribution of putative novel modifications. **f,** Upper, functional annotations of transcripts containing modifications in each sample type. Cell color corresponds to the highest-level annotation for each gene as defined in the Kyoto Encyclopedia of Genes and Genomes (KEGG): Genetic Information Processing (blue), Metabolism (purple), Cell Processes (gold), or Environmental Information Processing (green). Genes that did not have annotations in the KEGG database and those that were annotated as “general function” or “unknown function” were classified here as “Function unknown” (teal). Lower, identities of transcripts containing modifications. Cell size corresponds to the number of putative modified sites or regions in the transcript.

Putative modification regions were identified using two biological replicate samples per condition and two approaches for modification detection (**Fig. 2b; Extended Data Fig. 2a; Table S2)**. There was a low degree of overlap in putative modification regions between replicates and modification detection approaches, at 1.6% of sites in the no-stress samples **(Extended Data Fig. 2b–d)**. A similar phenomenon was observed in the inter-condition comparison, with just 1.2% of modifications found across all three conditions (**Fig. 2c**). mRNA modifications were largely found in the CDS and concentrated around the start and stop codons, independent of nucleotide identity or sample condition (**Fig. 2d, e**). Approximately 9.5% of the putative modification regions contained a previously identified pseudouridine modification motif^43^ within +/- 4 nt. Most of the modified mRNAs were involved in central functions, such as translation, transcription, central metabolism, and transport (**Fig. 2f**), indicating that stress-response genes were not primarily regulated at the epitranscriptomic level during the alarm phase.

### rRNAs undergo consistent epitranscriptomic changes across stress conditions

rRNA modification sites have previously been characterized in *E. coli*; however, stress-induced changes in modification abundance remain unclear. Using an IVT sample to control for systemic noise, we analyzed the oxidative-, acid-, and no-stress samples using the proportion of basecalling error (BCError) at each rRNA position to detect modification presence/absence. This approach allowed detection of 92% of the 36 known modified positions in the 23S and 16S **(Extended Data Fig. 3a, b)**. Because BCError can be used as a proxy for modification abundance^13,30–33^, we calculated normalized changes in rRNA modification levels between stress-treated (BCError_stress_) and no-stress (BCError_no stress_) samples as follows:

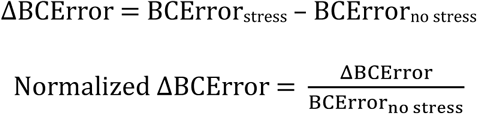

The 16S rRNA showed increased m^4^Cm1402 and m^5^C1407 levels under both stress conditions (**Fig. 3b**), as confirmed with MS **(Extended Data Fig. 3a–h)**. An m^5^C Rol-LAMP assay was used to experimentally validate these results at the single-nucleotide level. Under acid-stress conditions, the results were consistent with the sequencing data, demonstrating an increase only at position m^5^C1407 **(Extended Data Fig. 3i)**. Other changes (e.g., an increase in m^6,6^A at position 1519 of the 16S) were observed with sequencing but not confirmed by MS.

**Figure 3.**
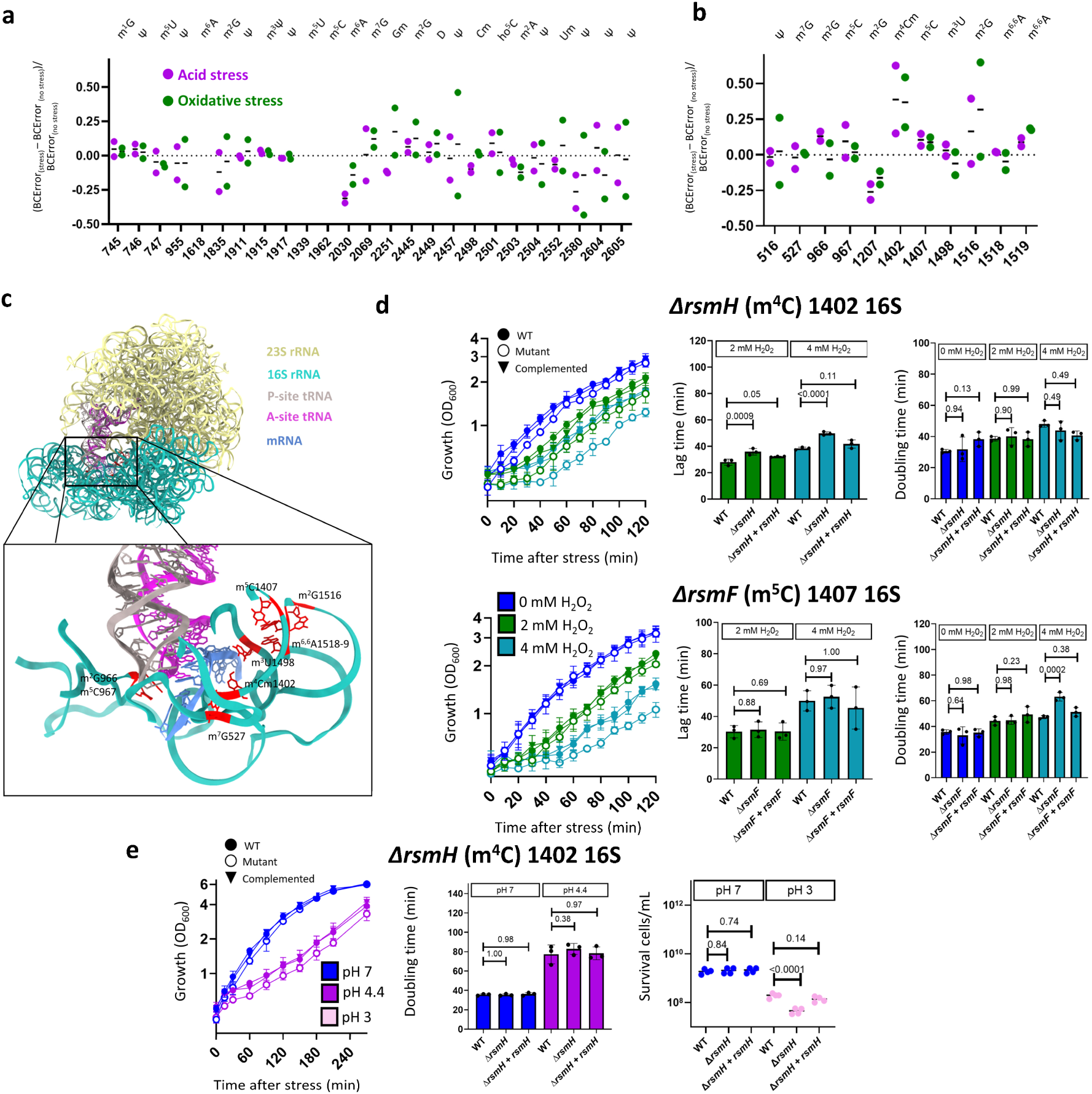
Epitranscriptomic changes in rRNA. a, b, Relative changes in **(a)** 23S and **(b)** 16S rRNA modification levels in acid- and oxidative-stress samples compared to the no-stress control. Values for each position were calculated as ΔBCError (BCError in the stress sample minus BCError in the no-stress control) divided by BCError in the no-stress control. **c,** Schematic showing the P and A sites of the ribosome with RNA modifications around the mRNA indicated (PDB 7k00). **d,** Phenotypic analyses of wild-type (WT) *E. coli* and Δ*rsmH* (m⁴C1402) and Δ*rsmF* (m⁵C1407) mutants under oxidative stress (2 or 4 mM H₂O₂), comprising growth rate (left), lag time (center), and doubling time (right). **e,** Phenotypic analyses of WT and Δ*rsmH* (m⁴C 1402) *E. coli* under acid stress (pH 4.4), comprising growth rate (left), doubling time (center), and survival (right). *p*-values calculated with two-way analysis of variance (ANOVA) and post-hoc Dunnett’s multiple comparisons test.

Notably, modifications with stress-induced changes were located near the ribosomal decoding center (**Fig. 3c**). To assess whether these modifications were required for stress responses, we generated knockout mutants for *rsmI* and *rsmH* (which are responsible for m⁴Cm1402), *rsmF* (m⁵C1407), *rsmJ* (m²G1516), and *rsmA* (m⁶,⁶A1518–1519), and compared growth phenotypes at 2 and 4 mM H_2_O_2_ (**Fig. 3d, Extended Data Fig. 4)**. Oxidative stress induced a longer lag phase in Δ*rsmH* and slower growth in Δ*rsmF* compared to the wild type (WT) (**Fig. 3d**). These results suggested that the corresponding modifications (m⁴C1402 and/or m⁵C1407) had key roles in the *E. coli* oxidative stress alarm phase. In contrast, the Δ*rsmI*, Δ*rsmJ*, and Δ*rsmA* knockouts showed no significant differences in growth compared to the WT under oxidative **(Extended Data Fig. 4a)**. Δ*rsmH* cells had a significant decrease in bacterial growth (i.e., biomass) and survival compared to the WT under acid stress (**Fig. 3e**). The Δ*rsmI*, Δ*rsmF*, and Δ*rsmA* knockouts showed no differences under acid stress **(Extended Data Fig. 4b–d)**. Overall, these results indicated that changes in rRNA modification levels were essential for overcoming growth inhibition (lag phase) during the alarm phase.

### Changes in the tRNA epitranscriptome are stress-specific

Changes in tRNA modification abundance were assessed using two sample types (**Fig. 2a**): mRNAe (described above) and Nano-tRNAseq (generated with a tRNA-specific library preparation kit^31^) **(Extended Data Fig. 5a)**. mRNAe samples were expected to contain a mixture of unprocessed, immature, and mature tRNAs, whereas the Nano-tRNAseq samples should be enriched in mature, aminoacylated (charged) tRNAs. Supporting this hypothesis, most Nano-tRNAseq reads were ∼70–100 nt in length (consistent with the mature tRNA size); mRNAe samples showed a much higher proportion of reads from 100–500 nt, corresponding to unprocessed and immature tRNAs (e.g., those that had not yet been cleaved of the leader and/or trailer sequences or other RNAs within the same operon)^8^ **(Extended Data Fig. 5b)**. Comparison of BCError between mRNAe and Nano-tRNAseq samples revealed higher average levels of each modification type in Nano-tRNAseq samples **(Extended Data Fig. 5c)**. Similarly, modification levels at each individual site across the tRNAs were generally higher in Nano-tRNAseq samples **(Extended Data Fig. 5d)**. Although there were no stress-dependent changes in tRNA expression **(Extended Data Fig. 5e, f)**, there were differences in relative tRNA abundance between the mRNAe and Nano-tRNAseq samples **(Extended Data Fig. 5g)**, further demonstrating that the two preparation methods captured distinct tRNA subpopulations.

To evaluate whether stress caused changes in tRNA modification levels, we compared BCError by calculating the difference between stressed samples and no-stress controls as previously described^30,31,34^. Stress exposure changed tRNA modification patterns only in the uncharged tRNAs from mRNAe samples (**Fig. 4a, b; Extended Data Fig. 6a–d)**, not in the charged tRNAs isolated with Nano-tRNAseq **(Extended Data Fig. 7a, b)**. Furthermore, MS was performed on a purified fraction of RNAs (≤ 200 nt) containing primarily mature tRNAs (comparable to those isolated with Nano-tRNAseq) to assess the general abundance of selected modifications. Very few differences in tRNA modification levels under oxidative and acid stress were found with MS **(Extended Data Fig. 8a–p)**, consistent with the Nano-tRNAseq sequencing data.

**Figure 4.**
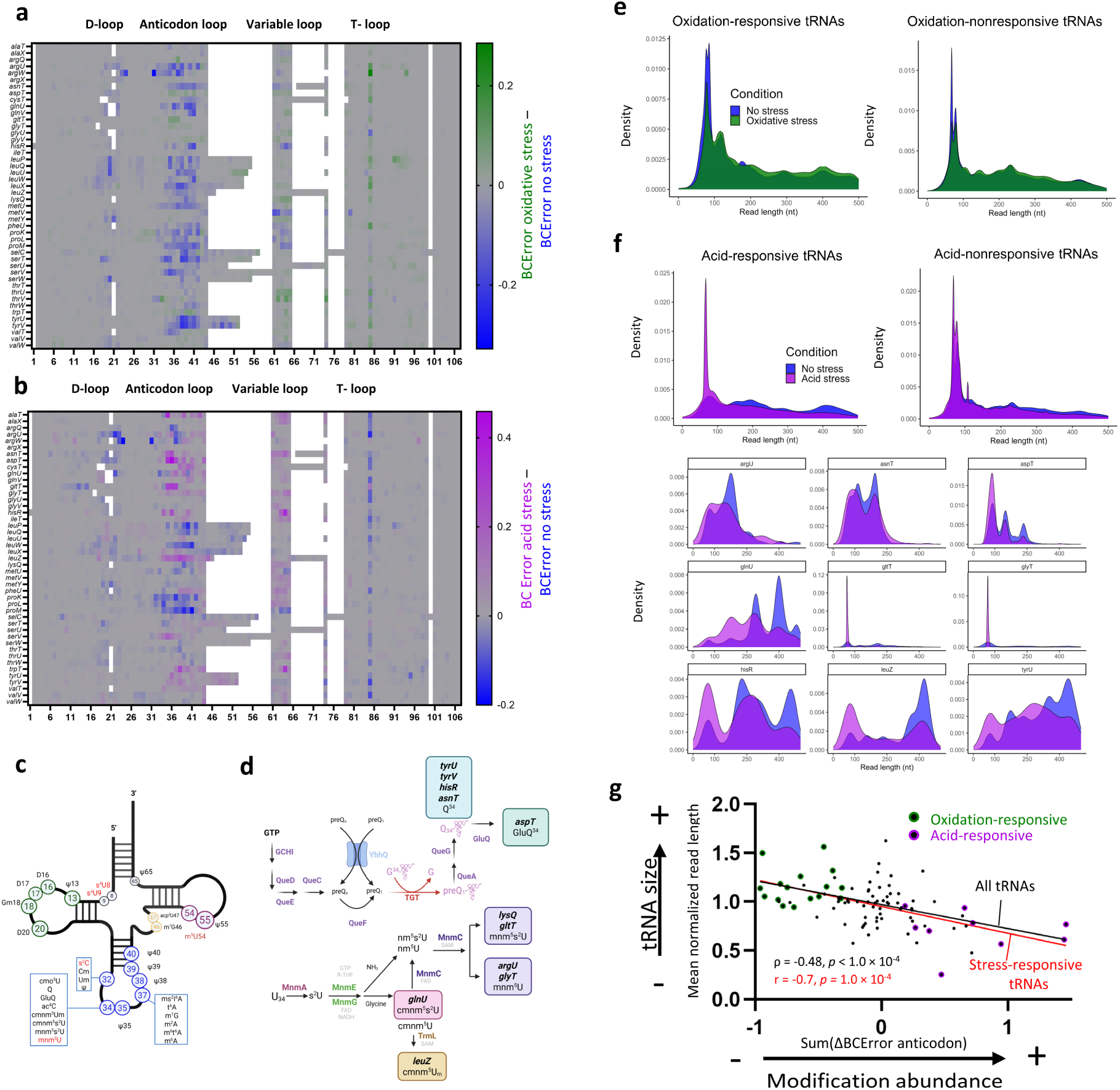
Epitranscriptomic changes in tRNA. a, b, Changes in uncharged *E. coli* tRNA modification abundance in response to **(a)** oxidative- and **(b)** acid-stress conditions. Changes in uncharged tRNA abundance at each tRNA position were calculated as BCError in the oxidative-stress sample minus BCError in the no-stress control for all **(a)** 46 and **(b)** 45 unique tRNAs present at a sufficient read depth (≥ 5 reads) and containing modifications that were detected with BCError. Data are shown as the average values from two biological replicates. **c,** Types and positions of characterized modifications in *E. coli* tRNAs. **d,** Schematic representations of the biosynthetic pathways from (upper) GTP to queuosine (Q) and (lower) UTP to Mnm. Q and Mnm are incorporated into five and six tRNAs, respectively. **e, f,** Normalized tRNA abundance at read lengths from 0–500 nt. Distributions are shown for **(e)** all tRNA species under no-stress and oxidative-stress conditions, separated by oxidative-stress responsiveness, and **(f)** each acid-responsive tRNA under no-stress and acid-stress conditions individually (upper) and separated by acid responsiveness (lower). Abundance was normalized to give equal weight to each tRNA, biological replicate, and sample condition regardless of read depth. **g,** Relationships between tRNA read length and anticodon loop modification abundance. The values for all tRNAs were non-normally distributed, whereas the values for the stress-responsive tRNAs alone were normally distributed (as determined with a Shapiro–Wilk test). The strengths of the relationships for all tRNAs (black) and stress-responsive tRNAs only (red) were therefore assessed with Spearman’s and Pearson correlations, respectively.

In mRNAe samples, oxidative stress led to a pronounced decrease in tRNA modifications within the anticodon loop of 16 tRNAs (**Fig. 4a**), which were thus considered oxidative-stress responsive. There were no significant differences in tRNA expression between control and stress conditions **(Extended Data Fig. 5e, f)**, indicating that stress-dependent changes were solely at the epitranscriptomic rather than the transcriptomic level. Acid stress produced a distinct response, with nine tRNAs exhibiting increased modifications at the wobble position (nucleotide 34) (**Fig. 4b, c**). Notably, all nine of these acid-responsive tRNAs have wobble-position modifications that are generated via the queuosine (Q) pathway^35^ (*aspT, asnT, hisR,* and *tyrU*) (**Fig. 4d, Extended Data Fig. 9a)** or the Mnm pathway^36,37^ (*argU*, *glnU, glyT, gltT,* and *leuZ*) (**Fig. 4d, Extended Data Fig. 9b)**.

To further explore the observed changes in tRNA modification levels, we analyzed the tRNA size distribution in stress-responsive compared to stress-nonresponsive tRNAs from mRNAe samples (**Fig. 4e, f**). Surprisingly, the oxidative-stress-responsive tRNAs were enriched in immature molecules under oxidative-stress compared to no-stress conditions, whereas non-responsive tRNAs showed comparable distributions under oxidative-stress and no-stress conditions (**Fig. 4e**). In contrast, the acid-stress-responsive tRNAs showed an accumulation of mature molecules under acid stress (**Fig. 4f**). This suggested that altered anticodon modification levels were associated with stress-induced variations in tRNA maturation. We therefore assessed the relationship between the average length of each tRNA under acid-stress compared to no-stress conditions (as a proxy for tRNA processing stage) and the cumulative ΔBCError for aligned tRNA positions 30–41 (as shown in **Fig. 4a, b**) (as a comprehensive measure of modification levels). There was a moderate correlation (ρ = -0.48, *p* < 1.0 × 10^-4^) among all tRNAs, which increased to a strong correlation (r = -0.70, *p* = 1.0 × 10^-4^) when only stress-responsive tRNAs were considered (**Fig 4g**). These data supported our hypothesis of stress-induced regulation of tRNA maturation during the alarm phase.

### Wobble-position modifications regulate translation of a subset of proteins under stress

The decoding process uses both Watson–Crick and non-Watson–Crick interactions for codon– anticodon pairing; the former does not require modifications in the anticodon for correct codon– anticodon interactions, whereas the latter are anticodon modification dependent. The observed increase in wobble-position modifications via the Q and Mnm pathways may indicate the presence of a stress-resistance mechanism via differential abundance of modification-dependent and - independent codons. To evaluate the effects of anticodon modifications on acid stress survival, we assessed the relationship between translational efficiency (TE) under acid stress^28^ and modification-dependent codon abundance. Notably, transcripts with increased TE at pH 4.4 vs. 7.6 had a higher abundance of modification-dependent codons than transcripts with decreased TE **(Extended Data Fig. 9c)**. This effect was not observed at pH 5.8 **(Extended Data Fig. 9d)**, indicating that the mechanism was only in operation under severe acid stress.

We next ranked all *E. coli* transcripts by the relative abundance of modification-dependent codons (see **Methods**) to determine whether MoTTs regulation may occur under acid stress. The 400 transcripts with the highest abundance of Q- and Mnm-pathway modification-dependent codons **(Table S3)** were enriched in functions associated with membrane transport, transcription, stress, and mobility (**Fig. 5a**). Conversely, the ∼400 transcripts with the lowest abundance of Q- and Mnm-pathway modification-dependent codons **(Table S3)** were generally related to central carbon metabolism, translation (e.g., ribosomal proteins), and transcription (e.g., RNA polymerase) (**Fig. 5a**). This enrichment of stress-responsive functions among transcripts with a high abundance of modification-dependent codons supported the idea that MoTTs regulation is involved in the alarm phase.

**Figure 5.**
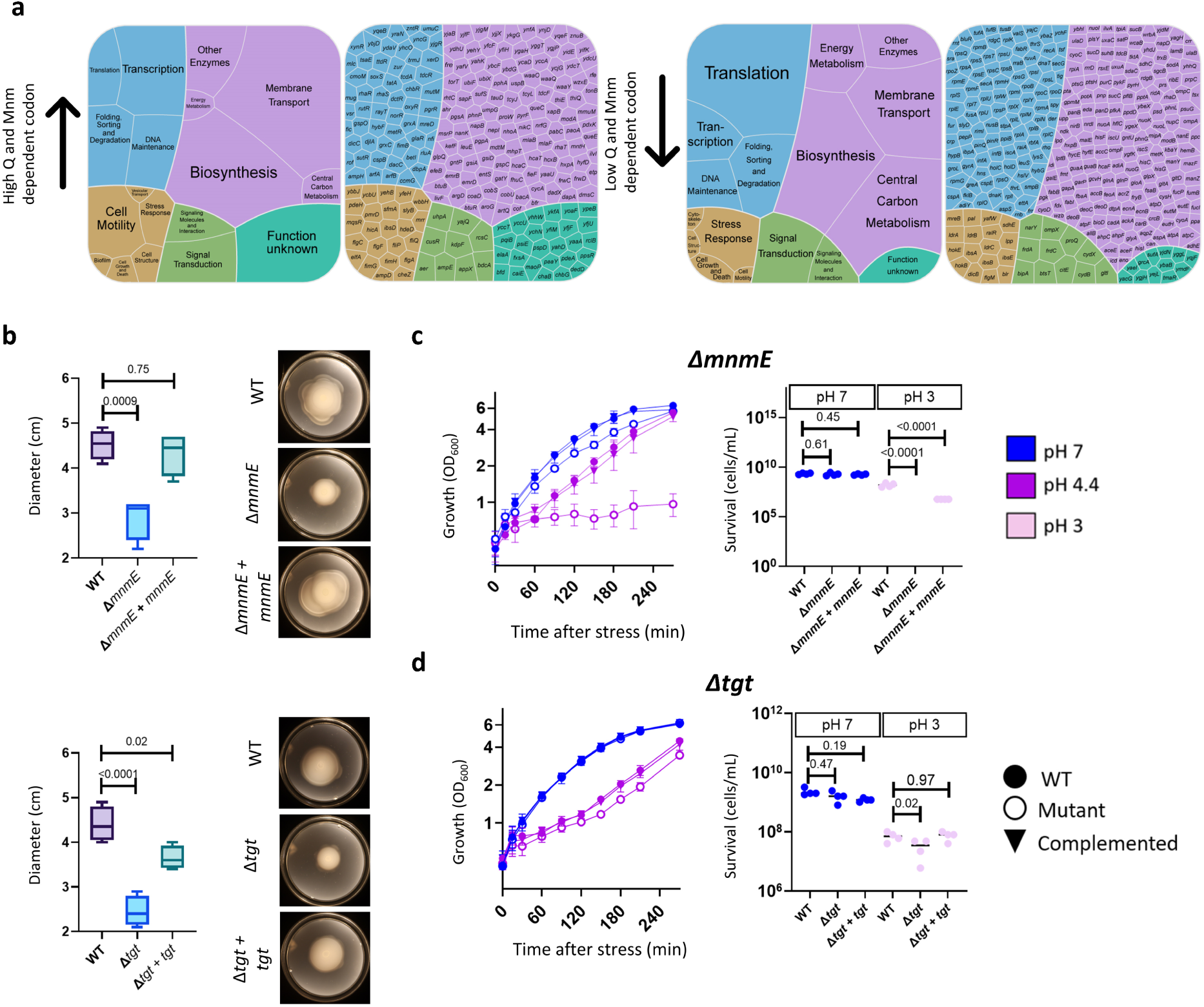
Modifications in the wobble position are implicated in *E. coli* stress recovery. a, Functional annotations of the 400 transcripts with the highest (left) and lowest (right) relative abundance of Q- and Mnm-dependent codons. **b,** Swimming motility of Δ*mnmE* (Mnm-pathway) and Δ*tgt* (Q-pathway) mutants compared to the WT. Data are shown as the average of four biological replicates. *p*-values calculated with one-way ANOVA and post-hoc Dunnett’s multiple comparisons test. **c, d,** Left, phenotypic analysis of **(c)** Δ*mnmE* and **(d)** mutants under acid stress (pH 4.4) compared to no-stress conditions. Right, cell survival after acid shock assays. Differences between strains were assessed using two-way ANOVA with post-hoc Dunnett’s multiple comparisons test.

One of the most specific functional annotations among transcripts with a high abundance of Q- and Mnm-pathway modification-dependent transcripts was cell motility. To investigate whether tRNA modifications enabling translation of these transcripts were necessary for motility, we phenotypically characterized mutants lacking enzymes required for wobble-position modifications via the Q and Mnm pathways. Specifically, we deleted genes encoding Tgt, an RNA-guanine transglycosylase responsible for Q modification^38^; MnmE, which is involved in mnm⁵s²U biosynthesis^39^; and MnmA, which catalyzes 2-thiouridine formation^40^ (**Fig. 4d**). These mutants showed impaired swimming capacity compared to the WT (**Fig. 5b, Extended Data Fig. 9e)**, demonstrating the essentiality of tRNA modifications in cell motility. Furthermore, Δ*mnmE* mutants displayed a strong growth defect and decreased survival compared to the WT under acid stress (**Fig. 5c**). The Δ*mnmA* mutant showed a similar phenotype **(Extended Data Fig. 9e, f)**, and although the Δ*tgt* mutant displayed only a minor growth defect at pH 4.4, its survival was impaired at pH 3 (**Fig. 5d**). These findings confirmed the critical roles of the Q and Mnm pathways^36^ in the *E. coli* acid stress response.

## DISCUSSION

Bacteria survive harsh conditions by detecting environmental changes, then employing diverse adaptation strategies. These adaptations involve transcriptional, translational, and post-translational regulation to alter gene expression and cellular metabolism, often mediated by sophisticated regulatory networks, global regulators, and small RNAs^41^. However, there has yet to be a comprehensive, systematic assessment of multiple stress responses at the epitranscriptomic level. To fill this gap, we used DRS to evaluate the epitranscriptomic profile during the alarm phase, an early stage of the stress response in which multi-level responses are initiated for survival and adaptation. The use of two abiotic stress conditions, acid and oxidative stress, allowed us to identify components of the alarm phase that are stress-specific and those that are more general.

For detection of putative novel modifications in the mRNA and ncRNA, we employed two approaches with distinct mechanisms: one involving evaluation of the BCError associated with each RNA position^30^, and one (nanoSundial) that uses raw electrical signal features from nanopore sequencing^42^. Overall, mRNA modifications were enriched in the CDS and found in transcripts associated with central functions, such as carbon metabolism and transcription. There was a high degree of variability between biological replicates, potentially explainable as off-target catalysis by rRNA- and tRNA-modification enzymes due to changes in substrate availability (a well-documented phenomenon in bacteria^13,43,44^; **Extended Data Fig. 10a**). However, in addition to prior studies showing that mRNA modifications can alter gene expression^43,45^, recent work has demonstrated strong conservation of specific mRNA modifications across bacterial phyla^40^. Taken together, these findings indicate important functional roles of bacterial mRNA modifications. Our set of putative novel modification sites offers an extensive pool of targets for future validation of such functions.

The rRNA epitranscriptome showed stress-type-independent changes during the alarm phase, which were phenotypically detectable as differences in the lag phase and growth rate. The increase we observed in m^5^C1407 of the 16S rRNA is consistent with previous findings that modification levels at this position increase under oxidative stress^46^. Furthermore, the changes observed in both m^4^Cm1402 and m^5^C1407 have also been detected under heat stress^13,46^, demonstrating that these modifications have roles in the alarm phase under multiple stressors. Heat stress leads to an increase in m^6,6^A1518– 19 levels^13^, but this effect was not observed here under oxidative or acid stress.

Notably, analysis of rRNAs via DRS, MS, and m^5^C Rol-LAMP under acid and oxidative stress revealed stress-dependent changes in modifications only within the rRNA decoding center, consistent with results under heat stress^13^. A recent study demonstrated that m^5^C1407 increases the duration of the excited and ground states, implying that this modification increases the available time for correct codon–anticodon recognition during translational elongation^47^. Thus, rRNA modifications in the decoding center may represent a dynamic control of the decoding process. This hypothesis is supported by our phenotypic mutant analysis, showing that RsmH and RsmF (which catalyze stress-responsive rRNA modifications) were important for both oxidative- and acid-stress responses. We therefore propose that increased modification levels in this region could decrease errors during decoding **(Extended Data Fig. 10b)**.

To thoroughly assess tRNA modifications during the alarm phase, we employed two distinct sample preparation methods. Multiple protocols have been developed specifically for tRNA sequencing, typically involving size-based fractionation, diacylation, and adapter ligation^30,31,34^ (e.g., Nano-tRNAseq). Alternative protocols rely on general polyadenylation (e.g., our mRNAe sample preparation method), which captures tRNAs together with other RNAs. The first method theoretically yields a higher read number, whereas the second allows for simultaneous sequencing of multiple RNA types within a single sample. Furthermore, the diacylation step required by approaches such as Nano-tRNAseq ensures that the captured tRNA population is enriched in fully mature (charged) tRNAs, in contrast to the mixed populations of immature, partially processed, and mature but uncharged tRNAs obtained with general polyadenylation. Importantly, our work is the first to report tRNA sequencing results from parallel implementation of polyadenylation^13,42^ and deacylation/primer ligation^30,31^ protocols.

Detected tRNA modification levels varied considerably between Nano-tRNAseq and mRNAe samples. The higher number and abundance of modifications detected in Nano-tRNAseq samples were consistent with previously published data indicating that tRNAs undergo sequential processing and modification to reach a form that is available for aminoacylation^8^. However, the mature, charged tRNAs captured with Nano-tRNAseq showed no statistically significant differences in modification levels under stress conditions compared to no-stress controls. In contrast, tRNAs from the mRNAe preparation showed a huge decrease in anticodon modifications under oxidative stress, consistent with effects previously observed under heat stress^13^. *E. coli* tRNA expression levels are greatly decreased at 5 min after H_2_O_2_ exposure, diminishing translation elongation^48^. Our results therefore suggest that the tRNAs present in mRNAe samples were newly synthesized and immature at 15 min after oxidative stress **(Extended Data Fig. 10c)**, and decreases in translation elongation are likely explained by both lower tRNA expression and decreased modification levels.

Compared to oxidative stress, a distinct pattern was observed in tRNAs under acid stress; although there was a decrease in pseudouridine levels in the T-loop, some sites in the anticodon wobble position showed increased modification levels. These acid-responsive tRNAs, regulated by the Q and Mnm pathways, were enriched in mature molecules available for aminoacylation. Generally, transcripts with a higher TE under acid stress had a greater abundance of codons requiring acid-responsive anticodon modifications for translation compared to transcripts with decreased TE, implicating stress-induced MoTTs regulation.

Our study provides the first comprehensive description of a multifaceted epitranscriptomic response in *E. coli* during the alarm phase of two stress conditions. We uncover simultaneous epitranscriptomic changes in the mRNA, tRNA, and rRNA, revealing a spectrum of regulatory events that range from stress-type-independent signatures to stress-specific adaptations. These findings highlight epitranscriptomic regulation as a previously underappreciated layer of the bacterial stress response, offering new insights into how bacteria rapidly reprogram their physiology to survive environmental challenges. By generating the most comprehensive dataset of RNA modifications in a model prokaryote to date, this work establishes a foundation for understanding the presence, abundance, and functional impacts of epitranscriptomic changes during the earliest stage of stress adaptations.

## ACKNOWLEDGEMENTS

We thank Sabine Peschek and Anna Chubanova for kindly providing mutant strains and plasmids. Some of the illustrations were created in BioRender.com

## Author contributions

Sebastián Riquelme-Barrios (Conceptualization, Methodology, Investigation, Formal analysis, Visualization, Writing—original draft, Writing—review and editing), Siobhan A. Cusack (Conceptualization, Methodology, Formal analysis, Visualization, Software, Data curation, Writing— original draft, Writing—review and editing), Luis Rivera-Montero (Methodology, Investigation, Formal analysis, Writing—review and editing), Leonardo Vásquez-Camus (Investigation, Writing—review and editing), Korinna Burdack (Investigation, Writing—review and editing), Sophie Brameyer (Investigation, Writing—review and editing), Maximilian Berg (Investigation, Writing—review and editing), G. Nur Yeşiltaç-Tosun (Investigation, Writing—review and editing), Stefanie Kaiser (Conceptualization, Funding acquisition, Methodology, Supervision, Writing—original draft, Writing— review and editing), Pascal Giehr (Conceptualization, Funding acquisition, Supervision, Methodology, Data curation, Software, Formal analysis, Visualization, Writing—original draft, Writing—review and editing), and Kirsten Jung (Conceptualization, Funding acquisition, Supervision, Methodology, Writing—review and editing).

## METHODS

### Strains and growth conditions

For experiments conducted with the WT *E. coli* strain MG1655, bacteria were cultured in lysogeny broth (LB) with 200 rpm shaking at 37 °C to an optical density at 600 nm (OD_600_) of 0.5. No-stress control cells were kept for 30 min at 37 °C to an OD_600_ of ∼0.8–1. For oxidative stress, 3% H_2_O_2_ was added to a final concentration of 2 or 4 mM in the bacterial culture. For acid stress, 5 M HCl was added to reduce the pH of the growth medium from 7 to 5.8. After 15 min, additional HCl was added to reduce the pH to 4.4. For both stress conditions, bacteria were grown at 37 °C for an additional 15 min after treatment before sample collection^28^. Strains carrying plasmids were grown overnight with chloramphenicol; the antibiotic was omitted during the stress experiment.

The single mutants Δ*rsmH*, Δ*rsmI*, Δ*rsmA*, Δ*rsmJ*, Δ*mnmA*, Δ*mnmE*, and Δ*tgt* were generated via in-frame deletion as previously described^13^. Primers used for mutant construction are listed in **Table S4**. Δ*rsmF* mutant strains were generated for a prior publication^46^. For complementation of Δ*rsmF*, the gene was inserted back into the *E. coli* genome in the original position with in-frame replacement. For complementation of Δ*rsmH*, Δ*mnmA*, Δ*mnmE*, and Δ*tgt*, the corresponding gene and its native promoter region were inserted into the *pBAD33* plasmid via Gibson assembly after linearization using two primers to eliminate the arabinose promoter **(Table S4)**.

### Total RNA isolation, tRNA/rRNA depletion, and mRNA enrichment

RNA was isolated using the PCI protocol^50^ with some modifications^12^. To produce mRNAe samples for nanopore sequencing, the resulting total RNA was treated to remove tRNAs and deplete ribosomes as previously described^13^. Primers for RT-qPCR to verify ribosome depletion are shown in **Table S4**. For MS analysis, RNA samples were separated using a size-selection protocol following the manufacturer’s instructions (RNA Clean & Concentrator Kit, Zymo Research, Irvine, CA, USA). The resulting > 200 nt fraction was used to assess rRNA modifications and the ≤ 200 nt fraction was used to evaluate tRNA modifications. Sample purity and size were confirmed via electrophoresis using the 2100 Bioanalyzer (Agilent Technologies, Santa Clara, CA, USA).

### RT-qPCR

cDNA was reverse transcribed from purified RNA samples and RT was carried out for qPCR to quantify expression of *katG*, *cadB*, *rrlH* (23S rRNA), *rrsH* (16S rRNA), and *recA* using previously described methods^13^. RT-qPCR primer sequences are shown in **Table S4**.

### MS analysis of RNA modifications

Absolute quantitative analyses using isotope dilution MS were performed as previously described^13^ with the addition of N4-acetylcytidine (ac^4^C) and aminocarboxypropyluridine (acp^3^U) (Carbosynth, Staad, Switzerland); Q (provided by the Peter Dedon laboratory); and 5-methylaminomethyl-2-thiouridine (mnm^5^s^2^U) (provided by the Susanne Häußler laboratory).

### Nanopore sequencing

IVT RNA control samples were prepared from no-stress samples as previously described^13^, with the addition of tmRNA depletion using custom probes supplied by siTOOLS (Planegg, Germany) as the final step. For mRNA-enriched samples and IVT controls, library preparation was carried out following the manufacturer’s protocol (ONT #SQK-RNA004). DRS was conducted with the FLO-MIN004RA (RNA) platform (ONT, Oxford, UK) using one flow cell per sample.

### Nano-tRNAseq

Total RNA samples were extracted, purified, and treated with DNase as previously described^13^. A total of 10 µg of RNA was used to isolate the tRNA fraction (≤ 200 nt) through size-selection protocols following the manufacturer’s instructions (RNA Clean & Concentrator Kit, Zymo Research, Irvine, CA, USA). RNA quality and successful size selection were confirmed by electrophoresis using the 2100 Bioanalyzer (Agilent Technologies, Santa Clara, CA, USA). Subsequently, 500 ng of tRNA-enriched samples were used for library preparation following the Nano-tRNAseq protocol (Immagina Biotechnology, Trento, Italy), based on a previously described method^31^. The protocol includes a diacylation step followed by splint adapter hybridization and ligation. Barcoded adapters were then annealed and ligated, followed by RT. Six samples were pooled and prepared using the Nanopore library preparation kit SQK-RNA004 (ONT, Oxford, UK) for RNA ligation adapter attachment. Finally, the libraries were loaded onto a FLO-MIN004RA (RNA) flow cell and sequenced according to the manufacturer’s protocol.

### ONT data processing

Reads from ONT sequencing were basecalled using Dorado v5.0.0 (ONT, Oxford, UK). SeqKit^54^ v2.8.2 was used to calculate basic statistics, namely the average read length, N50 length, and the total read and base numbers. Average Q scores were determined with NanoPlot^55^ v1.43.0. mRNAe samples were aligned with Minimap2^56^ v2.26 to a modified version of the ensemble *E. coli* reference genome (Escherichia_coli_str_k_12_substr_mg1655_gca_000005845.ASM584v2) from which duplicate tRNA and rRNA genes had been removed. Nano-tRNAseq samples were aligned to a reference file containing only *E. coli* tRNA sequences. Mapped reads were filtered for quality with BamQVFilter (https://github.com/JMencius/BamQVFilter) v0.1.0 at a threshold of Q score > 9 for mRNAe samples and Q score > 5 for Nano-tRNAseq samples. SAMtools^57^ v1.22 was used to remove secondary and chimeric alignments and unmapped reads. The resulting quality-filtered, primary-mapped reads were classified into RNA biotypes (tRNA, rRNA, protein coding, nontranslating CDS, pseudogene, or ncRNA) using EnsemblBacteria annotations (GCA_000005845); all reads that mapped to the *ssrA* transcript were classified as transfer-messenger RNA (tmRNA). For tRNA read length analyses (**Fig. 4e, f; Extended Data Fig. 5b)**, reads classified as mapping to the tRNA were trimmed with Cutadapt v4.9^58^, then the trimmed reads were mapped and Q-score filtered as described above. Read counts were quantified using Salmon^59^ v1.10.1. For differentially expressed gene (DEG) analysis, read counts from Salmon were imported into R using the ‘tximport’ package^60^, and DEGs were identified using DESeq2 (v1.38.3)^61^. Significant DEGs were obtained by filtering the results based on *p*-value (≤ 0.05), *q*-value (≤ 0.05), and log_2_(fold change) (≥ 2). All programs were run with default parameters unless otherwise specified.

### Putative modification site analyses

#### Biotype annotations

Putative modification sites outside the tRNA and rRNA were classified as belonging to the CDS, 5′ UTR, CDS, 3′ UTR, intergenic region, or ncRNA based on annotations in the *E. coli* K-12 substr. MG1655 ASM584v2 reference genome (RefSeq #GCF_000005845.2) and published UTR genomic coordinates^29^.

#### Modification detection

Sites or regions containing a modification were identified using nanoSundial^42^ with default parameters. In addition, the BCError at every position of each transcript was calculated from BAM files using the “get_bcerror_freqs.py” script described by White et. al^30^ (https://github.com/rnabioco/tRNA004/blob/main/src/modifications/get_bcerror_freqs.py). The output “BCErrorFreq” values were used in further analyses. A method of classifying sites as modified or unmodified based on BCError values was established using a combined no-stress sample (comprising biological replicates 1 and 2) and the combined IVT sample. ΔBCError was calculated for each nucleotide position as BCError_no stress_ – BCError_IVT_. Precision and recall of all known modification sites in the rRNA were calculated using a range of ΔBCError thresholds, once including and once excluding regions that were not classified as modified by nanoSundial. Based on this analysis **(Extended Data Fig. 2a)**, a ΔBCError threshold of 0.02 was used for classification of sites in the mRNA and ncRNA as putative modifications in the no-stress, acid-stress, and oxidative-stress samples **(Table S2)**.

#### Modified region characterization

The relative positions of putative modifications within a transcript were calculated based on the transcript length and the lengths of the CDS, 5’ UTR, and 3’ UTR. Modification abundance across the transcript was then visualized with the R package ‘ggplot2’^62^. For functional annotation visualization, the custom Kyoto Encyclopedia of Genes and Genomes (KEGG) hierarchy used by Proteomaps^63^ (https://www.proteomaps.net/data/KO_hierarchy.tms) was modified to include an additional ∼1000 *E. coli* genes with KEGG functional annotations **(Table S6)**. Transcripts and the corresponding annotations were then visualized as Voronoi treemaps with the R package ‘WeightedTreemaps’^64^.

### Modification-dependent codon abundance

nine sets of codons were analyzed, each comprising two codons that encode the same amino acid and differ in nucleotide sequence only at the third position **(Table S3)**. In every set, one codon requires a wobble-position modification generated via the Q or Mnm pathway in the corresponding anticodon for translation (modification-dependent) and the other codon does not (modification-independent). The total number of all 16 codons was counted in every *E. coli* CDS using a codon usage tool (https://www.bioinformatics.org/sms2/codon_usage.html), then the normalized modification-dependent codon abundance was calculated for each as follows:

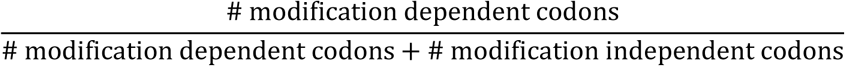

The 400 CDSs with the highest modification-dependent codon abundance **(Table S3)** were selected for further analysis. Due to a tie between the transcripts ranked in positions 399–403 for the lowest modification-dependent codon abundance, 403 transcripts were included in further analyses for this dataset **(Table S3)**. Functional annotations of the selected CDSs were visualized using Voronoi treemaps generated with the R package ‘WeightedTreemaps’^64^.

### Acid shock assay

Bacteria were grown in pH 7 LB medium at 37 °C to an OD₆₀₀ of 0.6–0.7. Cells were harvested via centrifugation for 2 min at 4000 × g, and 1 OD_600_ unit of cells were resuspended in LB at pH 5.8 or pH 7 as a control. After 15 min of growth at 37 °C, cells were harvested again and resuspended in pH 4.4 or pH 7 LB for another 15 min of growth at 37 °C. Cells were harvested one final time and resuspended in pH 3 or pH 7 LB, then grown for 1 h at 37 °C. After dilution in PBS, bacteria were plated on LB agar and grown overnight at 37 °C. Colony-forming units were then counted **(Extended Data Fig. 4c)**^65^.

### Motility assay

Bacterial cultures were grown overnight in LB broth at 37 °C. Cells were adjusted to an OD₆₀₀ of 1.0 in 1 mL of fresh LB media. Soft agar plates (1% tryptone, 0.25 g NaCl, and 0.3% agar) were prepared, and 10 µL of the cell suspension was spotted at the center of each plate. Plates were incubated for 12 h at 37 °C to allow bacterial growth and motility assessment. Colony size was quantified as the diameter.

### m⁵C-Rol-LAMP assay

m⁵C-Rol-LAMP was used to assess site-specific rRNA m^5^C methylation changes under pH stress^46^. Briefly, total RNA was treated for bisulfite conversion using the EZ RNA Methylation Kit (Zymo Research, Irvine, CA, USA); padlock probes were hybridized to the target sites and circularized by ligation **(Table S4)**. The circularized probes served as templates for rolling-circle amplification, followed by loop-mediated isothermal amplification. Reactions were performed with either dATP alone or dATP+dGTP. Signals were measured every 30 s (one measurement counted as one cycle) for 1 h using a CFX96™ Real-Time System (Bio-Rad, Hercules, CA, USA).

### Statistical analyses

Differences in the growth of mutant strains compared to the WT, in modification abundance between no-stress and acid-stress conditions (as quantified with m^5^C Rol-LAMP), and in *cadB* and *katG* expression (as quantified with RT-qPCR) were assessed with unpaired Student’s *t*-test. A ratio paired Student’s *t*-test was used to compare MS-based quantification of modification abundance under stress and no-stress conditions. Differences between mutant and WT cells in swimming motility and in lag time after acid shock were assessed with one-way analysis of variance with post-hoc Dunnett’s multiple comparison test. Two-way analysis of variance was used to determine statistically significant differences between mutants and WT strains with respect to doubling time and survival after acid shock; where this showed significant differences, a post-hoc Dunnett’s or Šidák correction was applied as appropriate. Differences in modification-dependent codon abundance between transcripts with significantly higher or lower translational efficiency under acid stress were assessed with a Mann– Whitney *U* test. The relationship between tRNA size and cumulative BCError in the anticodon was assessed with Spearman’s rank correlation and Pearson correlation in normally and non-normally distributed data, respectively, as determined with a Shapiro–Wilk test. An α value of 0.05 was applied for all statistical tests.

## Data availability

Sequencing data have been deposited at NCBI Gene Expression Omnibus (GEO) under accession number XXX.

This study does not describe any novel programs, software, or algorithms.

---

**Extended Data Fig. 1.**
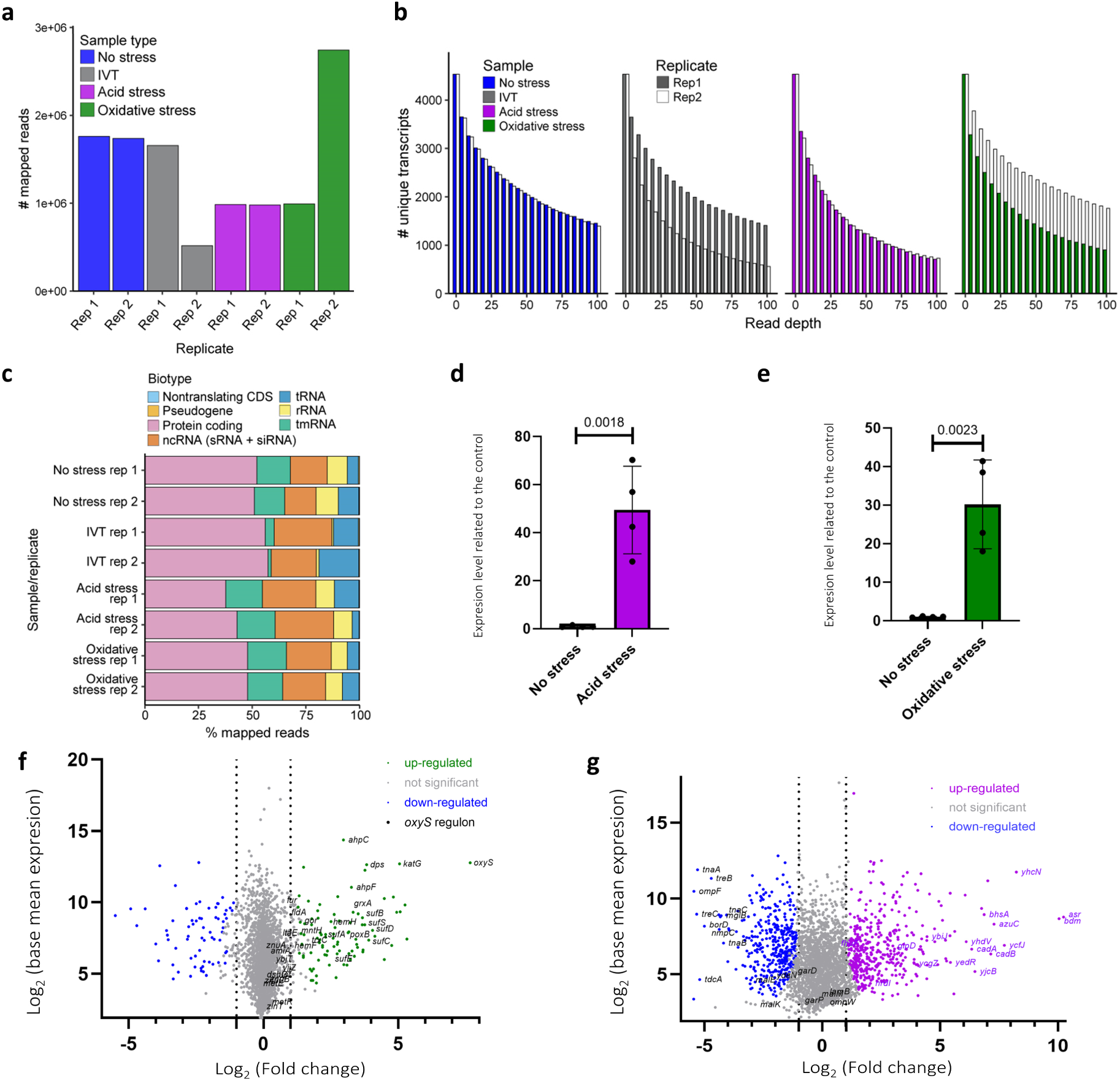
Sequencing quality analyses and differential gene expression in DRS data. a, Number of mapped reads produced from ONT sequencing of *E. coli* exposed to no stress, acid stress, or oxidative stress. Data are also shown for an *in vitro*-transcribed (IVT) control sample produced from the no-stress sample. **b,** Number of unique transcripts detected in the no-stress, IVT, acid-stress, and oxidative-stress samples across a range of minimum read depth thresholds. **c,** Relative abundance of reads mapped to distinct RNA biotypes in each sample. tRNA, transfer RNA; rRNA, ribosomal RNA; tmRNA, transfer-messenger RNA; ncRNA, noncoding RNA; sRNA, small RNA; siRNA, small interfering RNA. **d, e,** Expression of **(d)** *cadB*, which encodes a lysine/cadaverine antiporter, under acid stress and **(e)** *katG*, which encodes catalase/hydroperoxidase, under oxidative stress compared to the no-stress control. Expression was measured with quantitative reverse transcription quantitative PCR (RT-qPCR) and normalized to the 16S rRNA as an internal control. Significant differences were assessed with Student’s *t*-test. **f, g,** Differentially expressed genes (DEGs) in **(f)** oxidative- and **(g)** acid-stressed *E. coli* compared to a no-stress control. Data are displayed as log-transformed base mean expression vs. log_2_(fold change). In **f,** DEGs controlled by the *oxyS* regulon are labeled; in **g,** the 20 most strongly up- and down-regulated genes under acid stress conditions as previously described^28^ are indicated for comparison.

**Extended Data Fig. 2.**
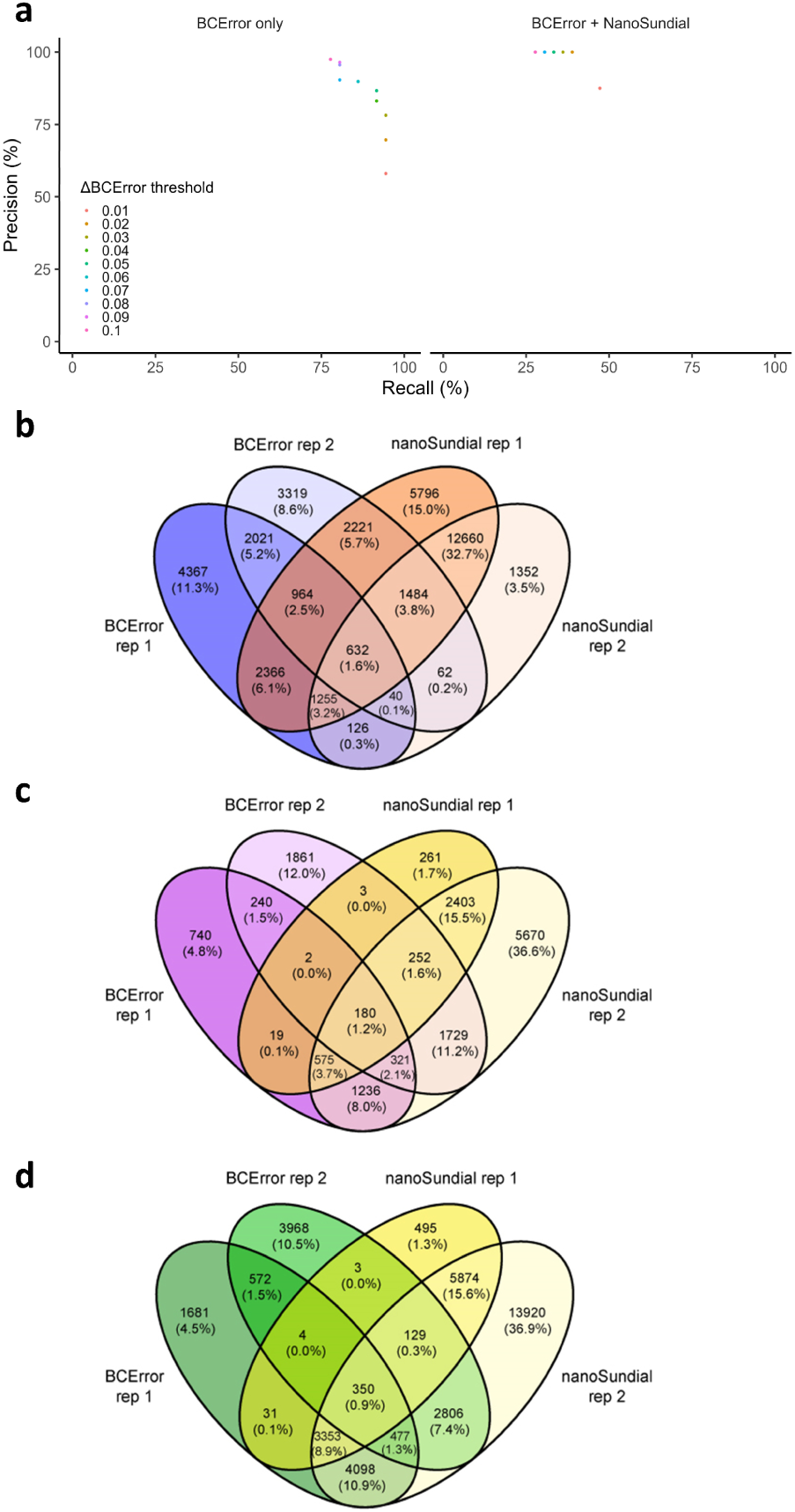
Detection of putative novel mRNA and ncRNA modifications in *E. coli* via DRS. a, Performance achieved using (left) BCError alone and (right) BCError plus nanoSundial to detect known *E. coli* rRNA modifications. **b–d,** Number of putative modification sites identified in the **(b)** no-stress, **(c)** acid-stress, and **(d)** oxidative-stress samples. Putative modification sites/regions were identified in two biological replicates of each sample type using two modification detection approaches, BCError and nanoSundial. The putative modification sites/regions were considered in further analyses only if they were present in both biological replicates and detected with both approaches.

**Extended Data Fig. 3.**
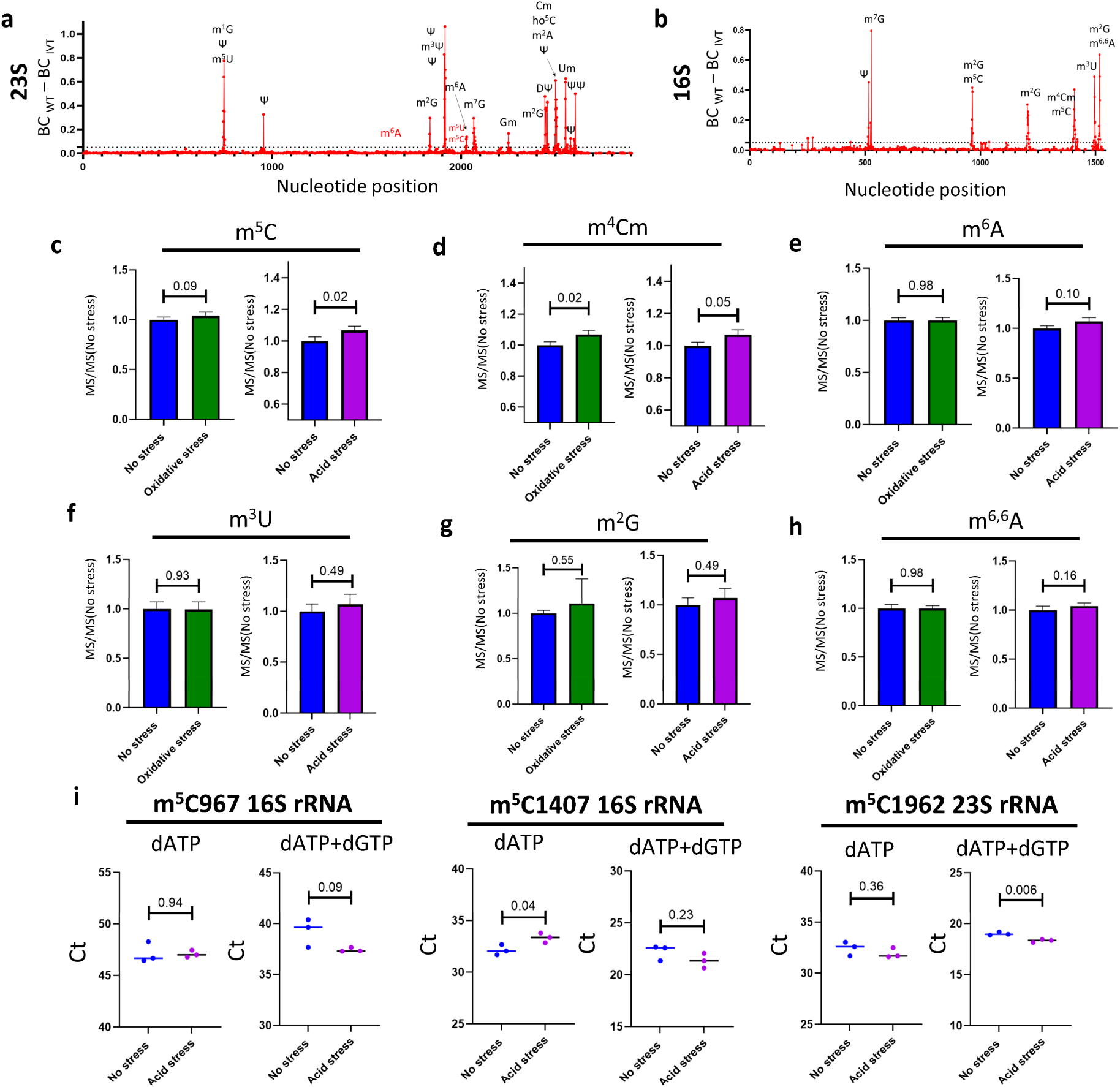
Orthogonal validation of changes in rRNA modifications. Detection of rRNA modifications in the **(a)** 23S and **(b)** 16S rRNA of *E. coli.* Values were calculated as the total percentage of basecalling error (BCError) at each site in no-stress samples minus the corresponding BCError value for each site in the IVT control. **c–h,** MS-based quantification of selected modification types in rRNA samples. Values were calculated as modification abundance in each sample divided by abundance in the no-stress control (MS/MS_No stress_) in four biological replicate samples. Error bars indicate the standard deviation of the mean. Significant differences were assessed with ratio paired Student’s *t*-test. **i,** Quantification of changes in m^5^C modification levels under acid stress for every reported m^5^C site in the rRNA via m^5^C Rol-LAMP. Total RNA was analyzed using the indicated site-specific padlock probes with dATP and dATP+dGTP in the ligation step. Data are shown for three biological replicates per sample type. Significant differences were evaluated with unpaired Student’s *t*-test.

**Extended Data Fig. 4.**
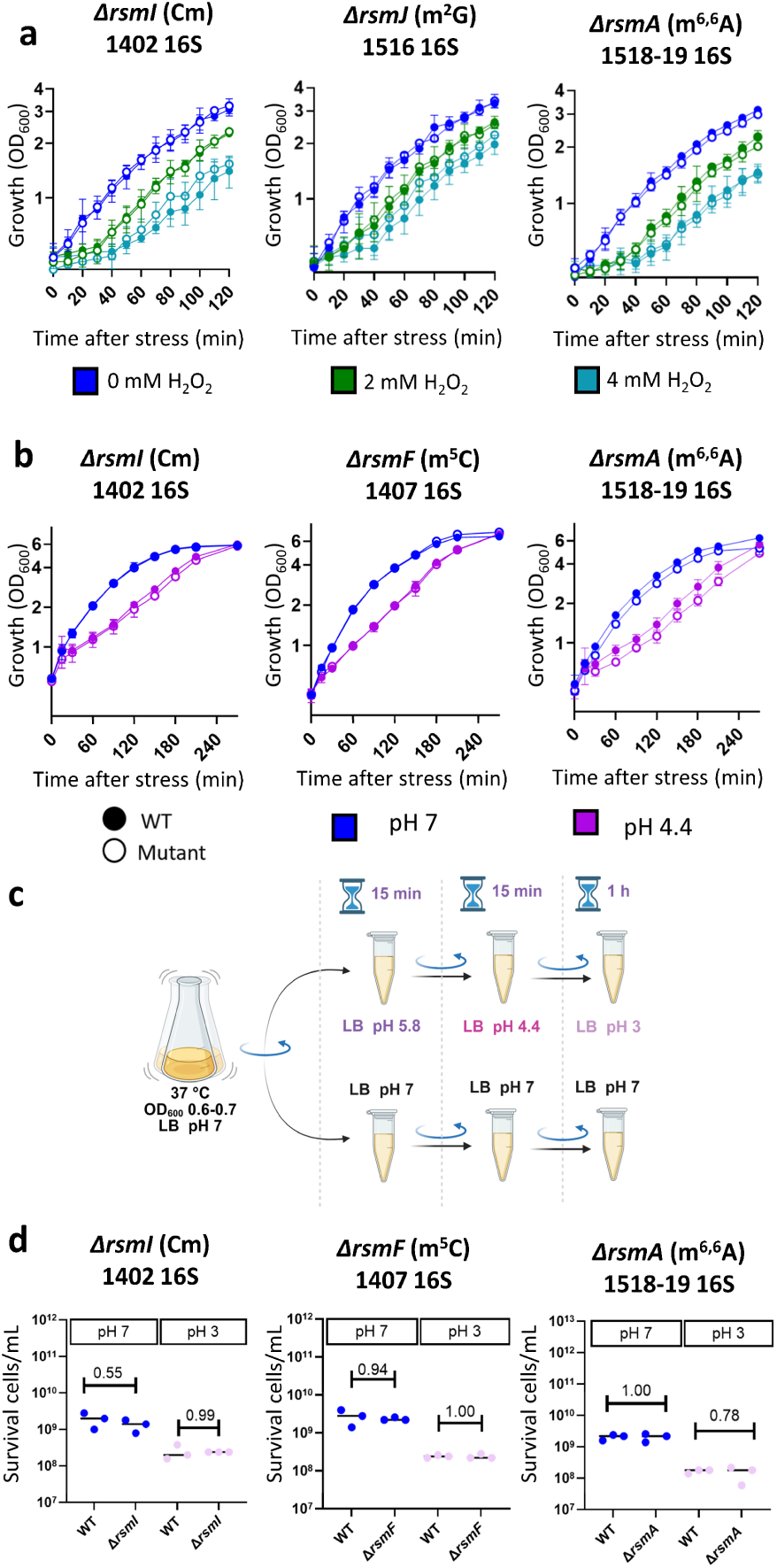
Phenotypic characterization of rRNA modification enzyme knockout mutants. a, Phenotypic analyses of WT *E. coli* and mutants that are unable to form known rRNA modifications, namely Δ*rsmI* (Cm1402 in the 16 rRNA), Δ*rsmJ* (m^2^G1516 in the 16S rRNA), and Δ*rsmA* (m^6,6^A1518–19 in the 16S rRNA). All strains were exposed to oxidative stress (2 or 4 mM H₂O₂). **b,** Phenotypic analysis of WT *E. coli* and the rRNA modification enzyme mutants Δ*rsmI* (Cm1402 in the 16 rRNA), Δ*rsmF* (m⁵C1407 in the 16S rRNA), and Δ*rsmA* (m^6,6^A1518–19 in the 16S rRNA) under acid stress. **c,** Scheme for acid shock assays (pH 3); briefly, cultures were grown at pH 7 or treated with a stepwise acid stress protocol (15 min at pH 5.8 followed by 15 min at pH 4.4) prior to exposure to LB medium at pH 3 for 1 h. Cultures were then serially diluted and plated on LB agar. **d,** Survival of WT, Δ*rsmI*, Δ*rsmF*, and Δ*rsmA* cells after acid shock treatment. *p*-values calculated with two-way ANOVA followed by Šidák correction for multiple comparisons.

**Extended Data Fig. 5.**
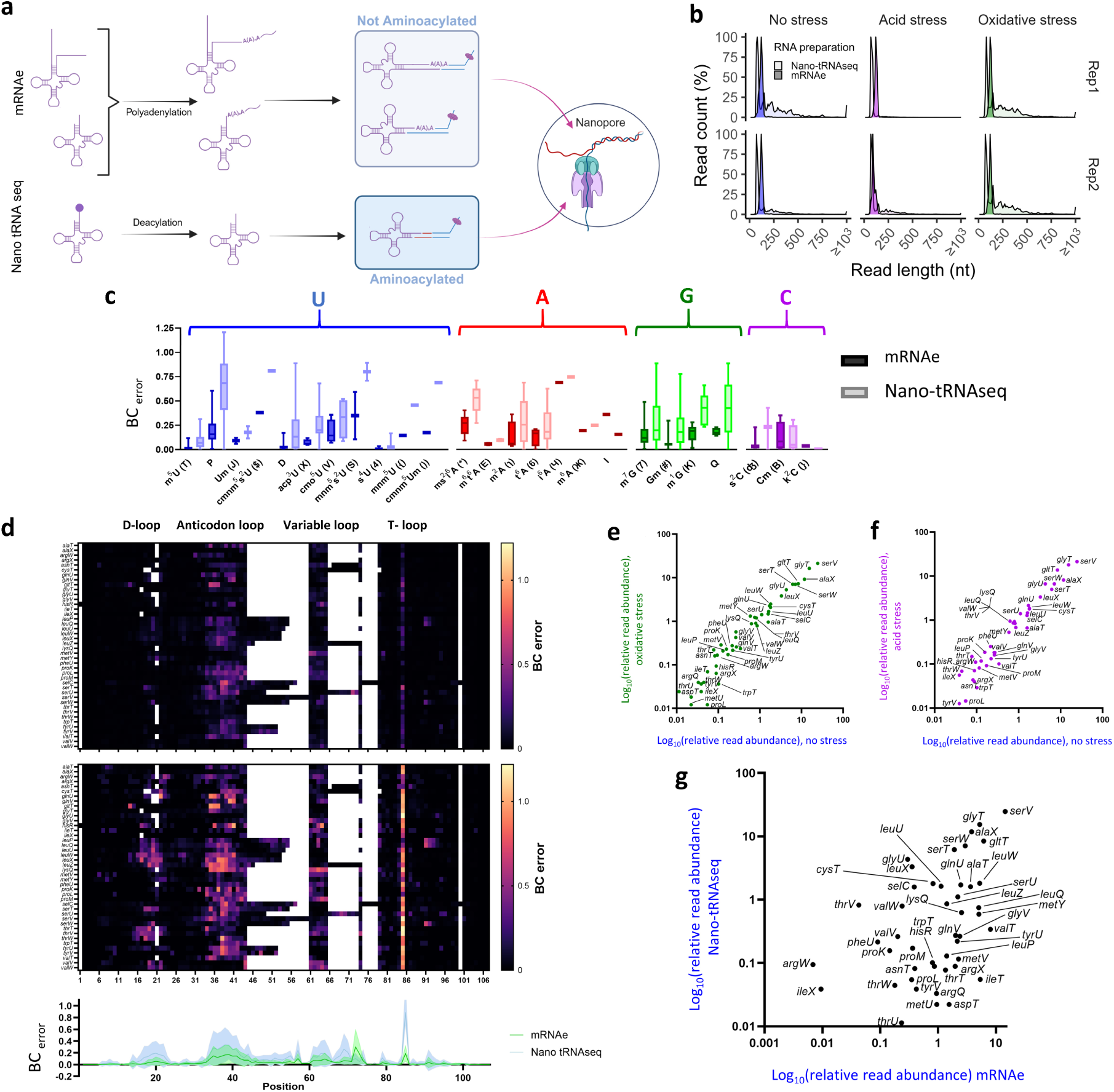
Nanopore DRS revealed distinct tRNA subpopulations. a, Graphical representation of the two protocols used for tRNA sequencing via nanopore. The mRNA-enriched (mRNAe) protocol captures uncharged tRNAs. This includes fully immature (unprocessed) tRNAs, which retain untranslated regions (UTRs) and/or lack the modifications required for aminoacylation; tRNAs at all stages of the maturation process; and fully processed, mature but uncharged tRNAs. In contrast, Nano-tRNAseq selects for aminoacylated tRNAs before library preparation, yielding samples that are highly enriched in mature tRNAs. **b,** Read length distribution for sequences mapping to the tRNA in two biological replicates of each sample type. Samples prepared with the Nano-tRNAseq protocol produced primarily reads of around 100 nt in length. Samples prepared with the mRNAe protocol produced the largest volume of tRNA reads around 90 nt, with a lower, broader peak from 90–500 nt. **c,** Comparison of BCError at sites with known modifications in no-stress samples prepared with the mRNAe and Nano-tRNAseq protocols. **d,** BCError at every nucleotide position of tRNAs detected in each sample type prepared with the mRNAe (upper) and Nano-tRNAseq (lower) protocols. Biological replicate samples were combined and a minimum depth of five reads was applied for inclusion in the analysis. **e, f,** Average relative abundance of reads mapped to each unique tRNA under **(e)** oxidative- or **(f)** acid-stress conditions compared to the no-stress control. **g,** Average relative abundance of reads mapped to each unique tRNA in no-stress samples prepared with the Nano-tRNAseq vs. mRNAe protocols. Significantly differentially abundant tRNAs between sample preparation methods were classified at *p* ≤ 0.05, *q* ≤ 0.05, and log_2_(fold change) ≥ 2.

**Extended Data Fig. 6.**
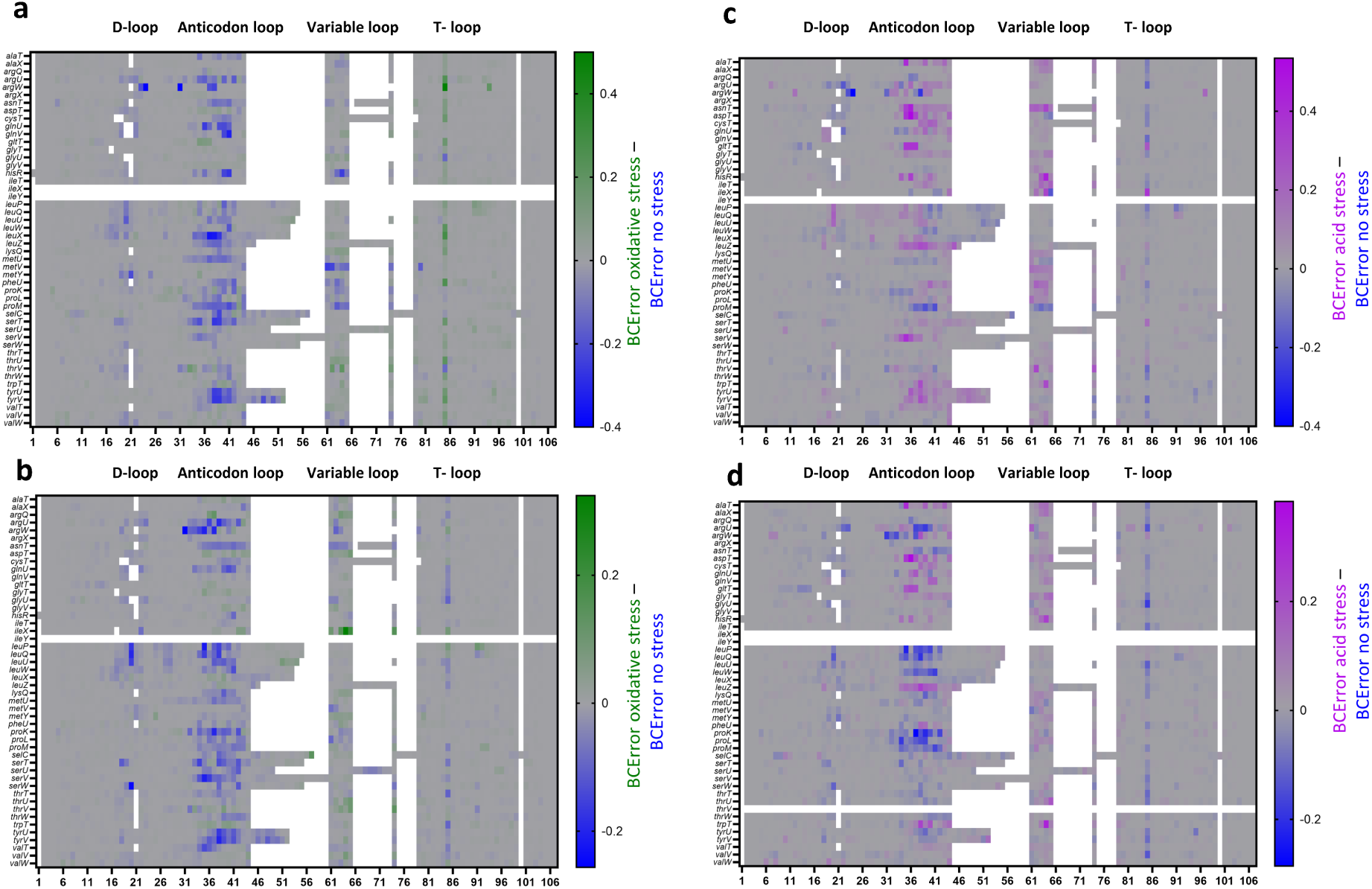
Single-sample changes in *E. coli* tRNA modification abundance during the stress alarm phase. tRNA abundance in biological replicates one (upper) and two (lower) of the **(a, b)** oxidative and **(c, d)** acid stress samples. tRNA was prepared and sequenced with the Nano-tRNAseq protocol; modification levels were determined with BCError. Values for each tRNA position were calculated as BCError_Stress_ – BCError_No stress_.

**Extended Data Fig. 7.**
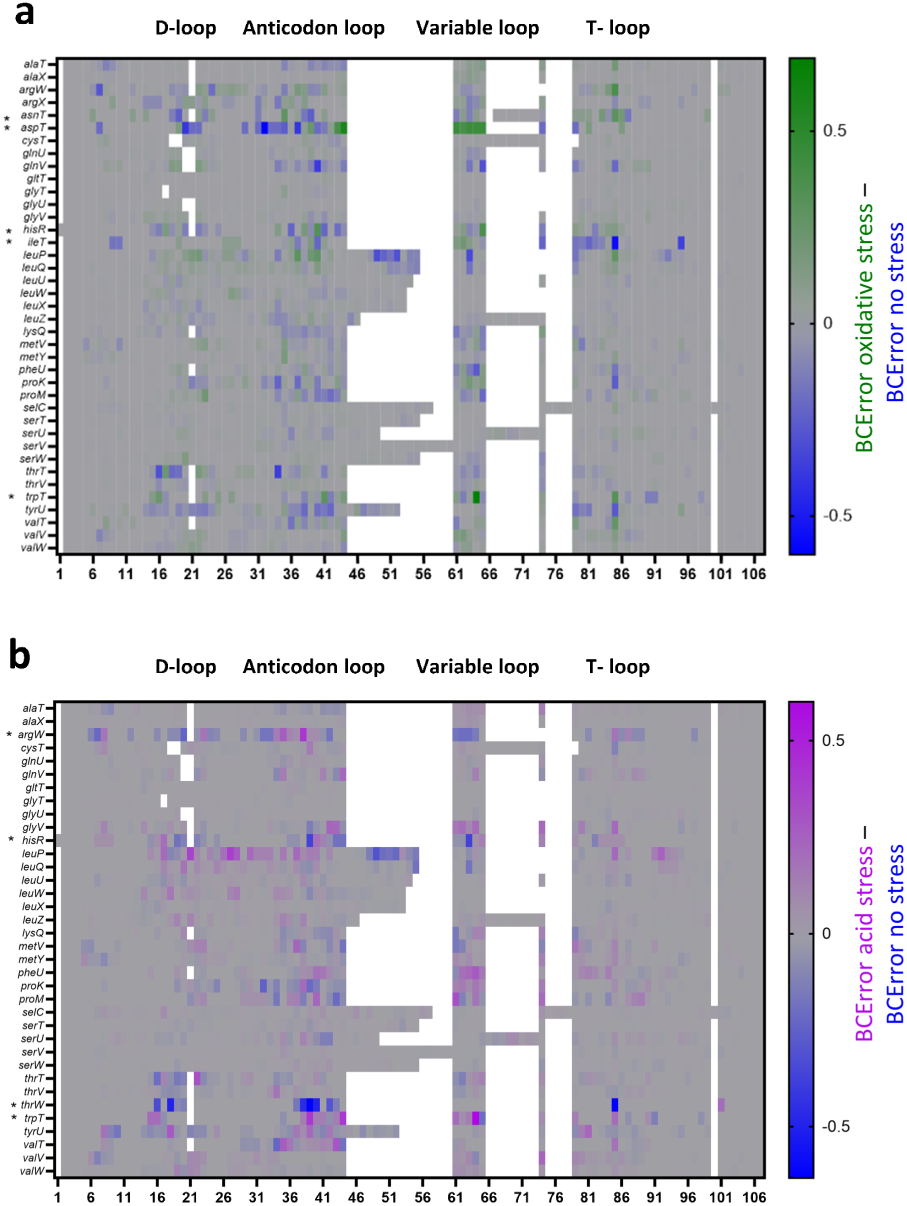
Average changes in *E. coli* tRNA modification abundance during the stress alarm phase. a, b, tRNA abundance levels in **(a)** oxidative-stress and **(b)** acid-stress samples prepared and sequenced with the Nano-tRNAseq protocol as determined with BCError. Values for each tRNA position were calculated as BCError_Stress_ – BCError_No stress_. Data are shown as the average of two biological replicates; asterisks indicate tRNAs that were present in only one biological replicate. tRNAs were classified as detected at a read depth of ≥ 5.

**Extended Data Fig. 8.**
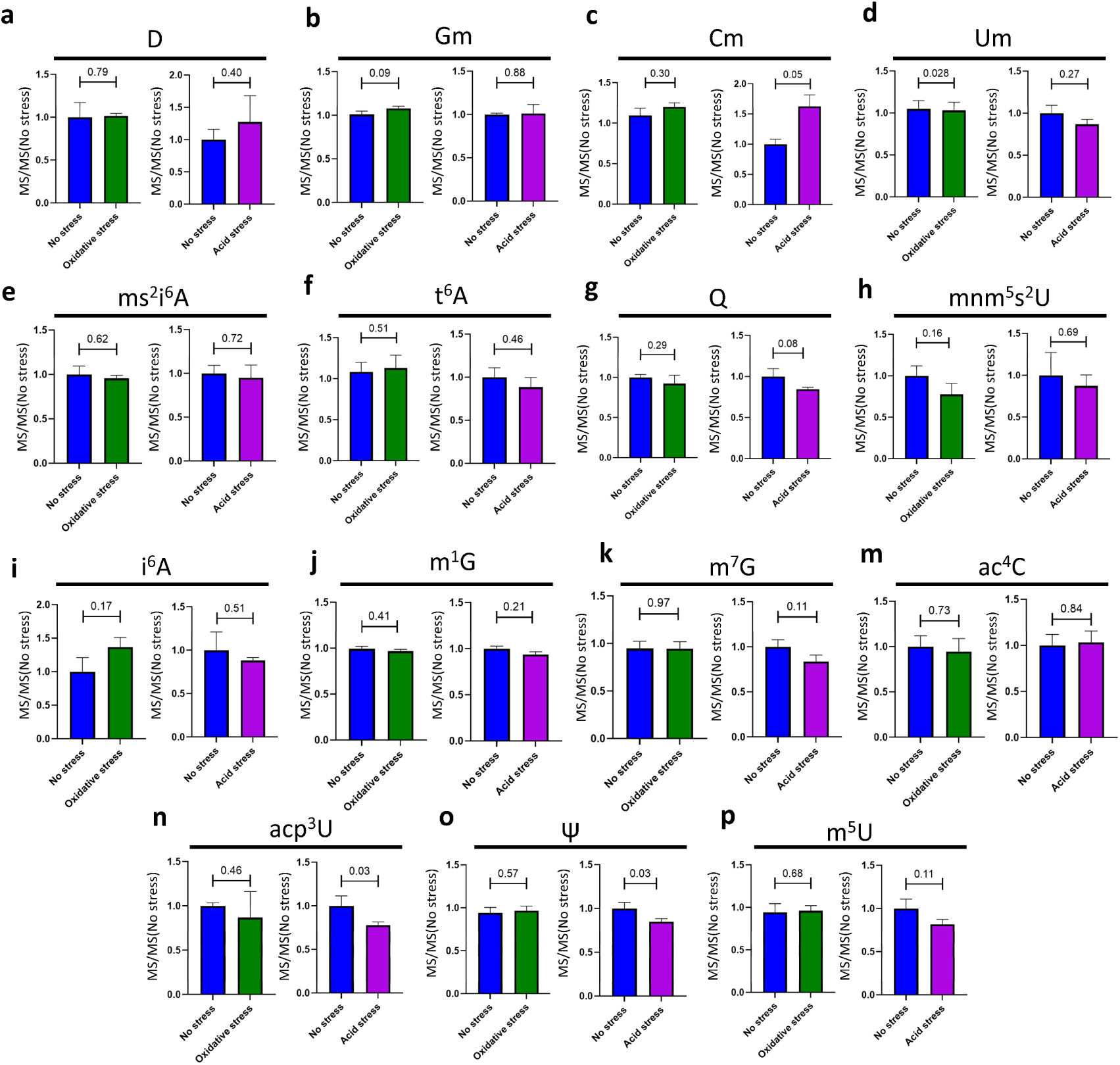
Mass spectrometric measurements of modifications in *E. coli* tRNAs. a–p, MS-based quantification of selected modifications in tRNA samples. Measurements were generated from purified RNAs of fewer than 200 nt in size. Modification abundance in each sample type was normalized to abundance in a no-stress control (MS/MS(No stress)). *p*-values calculated with ratio paired Student’s *t*-test.

**Extended Data Fig. 9.**
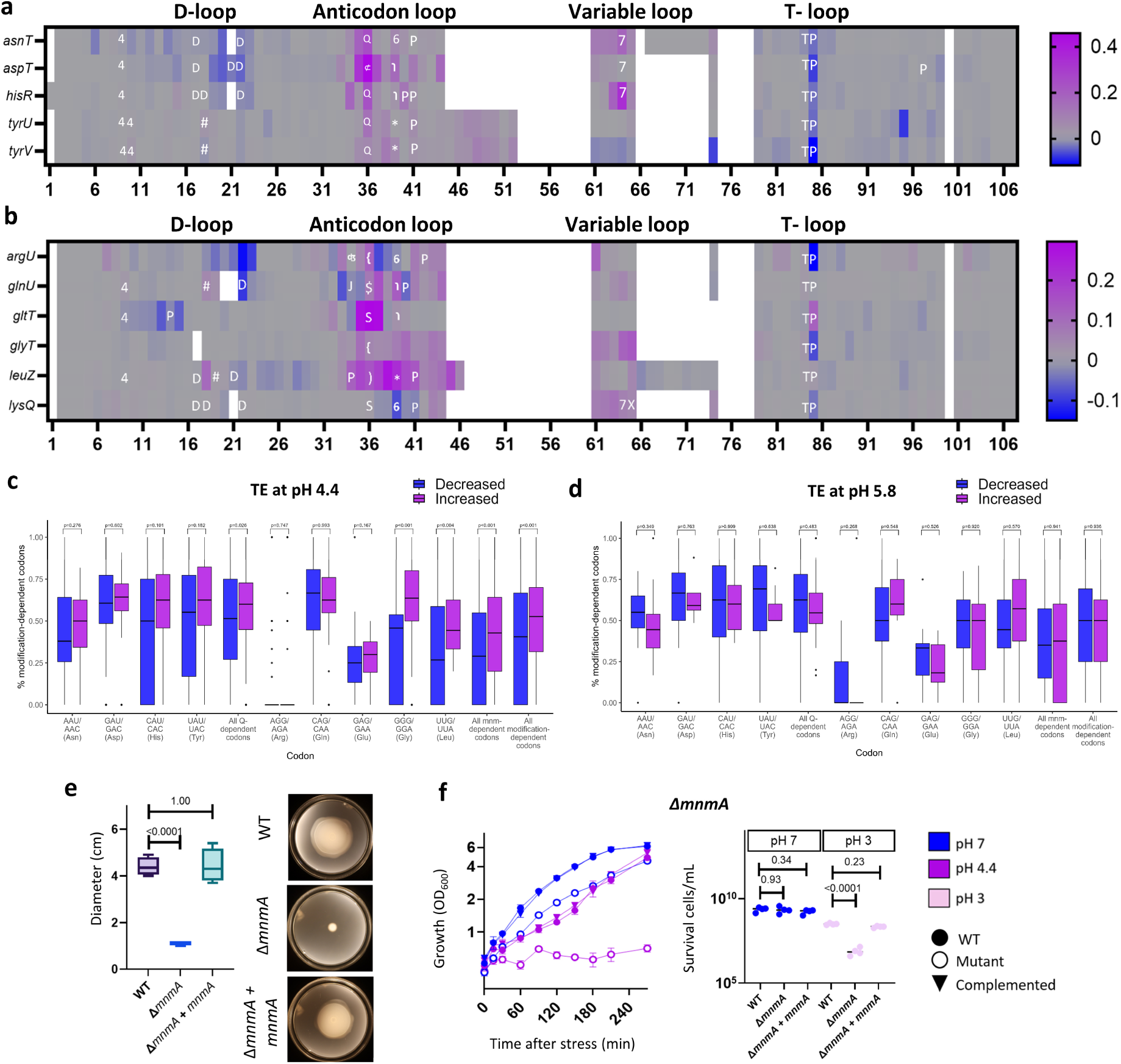
Δ*mnmA* mutants show similar phenotypes as Δ*mnmE* and *Δtgt* mutants. a, b, Changes in the abundance of **(a)** Q- and **(b)** Mnm-pathway modifications in tRNAs from mRNAe samples. Values were calculated as BCError_Acid stress_ – BCError_No stress_ for all positions across tRNAs containing Q or Mnm modifications. Data are shown as the average of two biological replicates. Symbols associated with the modification types are consistent with those used in the MODOMICS database^66^: ʤ, 2-thiocytidine (s^2^C) 4, 4-thiouridine (s^4^U); D, dihydrouridine (D); #, 2′-*O*-methylguanosine (Gm); ⊄, glutamyl-queuosine (gluQ); Q, queuosine (Q); *, 2-methylthio-*N*^6^-isopentenyladenosine (ms^2^i^6^A); ), 5-carboxymethylaminomethyl-2′-*O*-methyluridine (cmnm^5^Um); $, 5-carboxymethylaminomethyl-2-thiouridine (cmnm^5^s^2^U); S, 5-methylaminomethyl-2-thiouridine (mnm^5^s^2^U); {, 5-methylaminomethyluridine (mnm^5^U); 6, *N*^6^-threonylcarbamoyladenosine (t^6^A); P, pseudouridine (P); T, 5-methyluridine (m^5^U); ɿ, 2-methyladenosine (m^2^A); 7, 7-methylguanosine (m^7^G); X, 3-(3-amino-3-carboxypropyl)uridine (acp^3^U); J, 2′-*O*-methyluridine (Um). **c, d,** Proportion of codons requiring modifications synthesized via the Q or Mnm pathways for translation in transcripts having significantly increased or decreased translational efficiency at **(c)** pH 4.4 or **(d)** pH 5.8 compared to neutral pH according to previously published data^34^. **e,** Swimming motility of Δ*mnmA* mutant and WT *E. coli*. Data are presented as the average of four biological replicates. *p*-values calculated with one-way ANOVA and post-hoc Dunnett’s multiple comparisons test. **f,** Left, phenotypic analysis of WT and Δ*mnmA* mutants under no-stress and acid-stress (pH 4.4) conditions. Right, survival of WT and Δ*mnmA* mutants under acid stress (pH 3). Stress was induced by the addition of HCl to cultures at OD₆₀₀ = 0.5, then cells were monitored for 4.5 h. Significant differences in survival were assessed with two-way ANOVA and post-hoc Dunnett’s multiple comparisons test.

**Extended Data Fig. 10.**
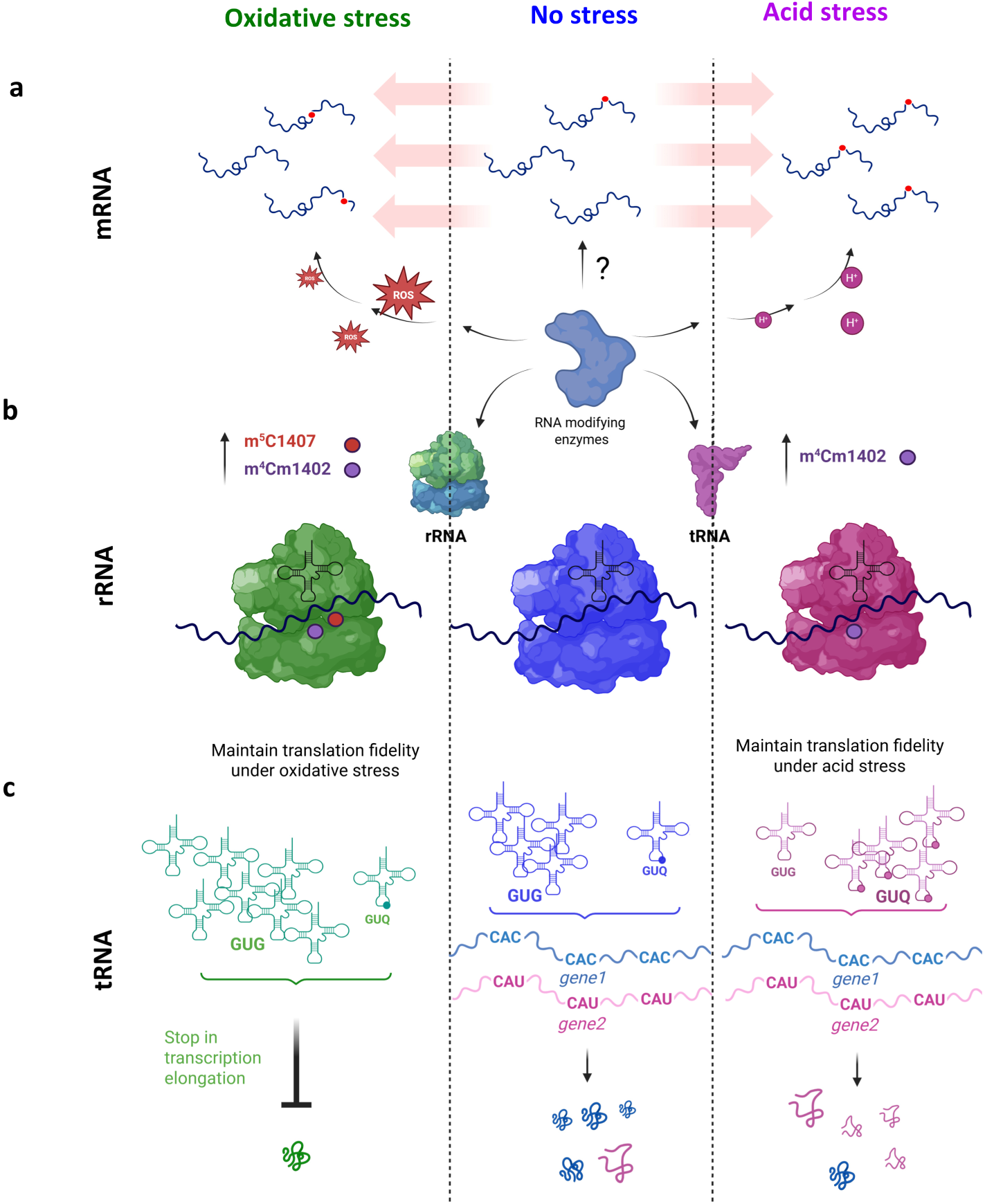
Working model of epitranscriptomic regulation under stress conditions in *E. coli*. a–c, Proposed models of **(a)** mRNA, **(b)** rRNA, and **(c)** tRNA modification levels in response to acid- and oxidative-stress conditions. Modification levels in mRNAs are generally low, whereas tRNAs and rRNAs are highly modified. Stress conditions such as acid or oxidative stress alter modification levels in the tRNAs and rRNAs to promote maintenance of essential cellular functions. Increased levels of the 16S rRNA modifications m⁴Cm1402 and m⁵C1407 in response to stress conditions may have a protective role, potentially maintaining translational fidelity during stress. Under acid stress, tRNA modifications may contribute to stress adaptation by enhancing the translation efficiency of stress-response proteins, independent of changes in RNA abundance.

**Table S1.** Basic statistics from ONT sequencing of no-stress, acid-stress, oxidative-stress, and IVT control samples.

**Table S2.** Genomic positions corresponding to regions classified as modified in the no-stress, acid-stress, and oxidative-stress samples. ΔBCError refers to the basecalling error frequency (BCError) at each position in the indicated sample with the BCError of the combined IVT sample at the same position subtracted. For regions (as opposed to single-nt sites) containing a putative modification, ΔBCError values are given for the site in that region with the highest BCError value.

**Table S3.** Relative abundance of Q- and Mnm-pathway dependent codons by transcript. The tab “Codons” shows the analyzed sets of modification-dependent and -independent codons. “400 most abundant” and “403 least abundant” show the ∼400 transcripts with the highest and lowest proportions of Q- and Mnm-dependent codons, respectively.

**Table S4.** Oligonucleotides used in this study.

**Table S5.** Mass transitions used in mass spectrometry analyses.

**Table S6.** Hierarchical KEGG annotations used for functional annotation analyses. The hierarchical classification scheme for *E. coli* used by Proteomaps^63^ was expanded to include an additional ∼1000 genes with annotations present in the KEGG database.

